# The evolutionary history and modern diversity of triterpenoid cyclases

**DOI:** 10.1101/2024.10.28.620730

**Authors:** Hanon Solomon McShea, Robb A. Viens, Babatunde O. Olagunju, José-Luis Giner, Paula V. Welander

**Affiliations:** Department of Earth System Science, Stanford University, Stanford, CA, USA; Department of Microbiology and Immunology, University of California, San Francisco, San Francisco, CA, USA; Department of Chemistry, SUNY-ESF, Syracuse, NY, USA

## Abstract

Cyclic terpenoids are a class of lipid compounds containing immense structural and functional diversity, with many cyclic triterpenoids acting as regulators of the physical properties and spatial organization of lipid membranes. Cyclic terpenoids are also readily preserved as terpane fossils, such as steranes and hopanes, forming a rich record of the evolution of life on Earth. Formation of the multiple ring structure of all cyclic terpenoids is catalyzed by terpenoid cyclase enzymes, among which are whole clades of proteins – many from environmental metagenomes and uncultured organisms – whose substrates and products are completely unknown. We investigate the function of these divergent cyclases through biochemical assays, and the evolutionary processes that produced them by testing and applying a variety of evolutionary models. We find deep divergence between the diterpenoid cyclases and triterpenoid cyclases, with other clades branching between the two, rooting the triterpenoid cyclase subtree, between squalene-hopene cyclases and sterol cyclases. Through a simple test of evolutionary rate shifts, we find an elevated evolutionary rate in the enzyme active site on the squalene-hopene cyclase stem, potentially indicative of positive selection. Finally, by testing the activity of divergent cyclases for a variety of substrates, we find a group of early-branching sterol cyclases from bacteria that synthesize arborinols, two of which produce the molecular precursor to a Permian “orphan biomarker.” Together, our data present an evolutionary framework for triterpenoid cyclases that can inform both the biochemical potential of these proteins and their products’ occurrence in the geological record.

## Introduction

In 1971, “geohopanes” were discovered in vast quantities in organic-rich sediments, ancient rocks, and fuel reservoirs (Ourisson & Albrecht, 1992). In the following years, hopanoid molecules from diverse bacteria were identified as the biological source of these molecular fossils (Förster et al., 1973; Ourisson & Rohmer, 1992). The finding precipitated a new era of research on the taxonomic distribution and physiological function of cyclic triterpenoids (hopanoids, sterols, and similar molecules shown in Figure 1A) and the triterpenoid cyclase enzymes that synthesize them, motivated by questions of what the molecular fossil record could reveal about the evolution of life on Earth, and of Earth itself (e.g. Fischer, 2008; Ourisson & Nakatani, 1994; Runnegar, 1991; Schaeffer et al., 1994). This body of research has uncovered diverse triterpane fossils, as well as an increasingly deep understanding of cyclic terpenoid structural diversity (Hayashi et al., 2007; Kontnik et al., 2008; Moosmann et al., 2020; Rudolf et al., 2021), taxonomic distribution (Mayer et al., 2021; Pearson et al., 2003; Ricci et al., 2014; Takishita et al., 2017; Villanueva et al., 2014, p. 201; Wei et al., 2016), biosynthesis (Banta et al., 2015; Brown et al., 2023; Lee et al., 2018, 2023; Pan et al., 2015; Pollier et al., 2019; Welander et al., 2010), and function (Brenac et al., 2019; Flesch & Rohmer, 1987; Gudde et al., 2019; Rivas-Marin et al., 2019; Schmerk et al., 2015; Welander et al., 2009, 2012; Welander & Summons, 2012; Zhai et al., 2024). Unsurprisingly, the catalogues of triterpane fossils and modern triterpenoids are not isomorphic – there are many “orphan biomarkers” in the geological record with no known biological source (Brocks & Pearson, 2005), and conversely, there are many organisms and environments that have genes for triterpenoid cyclases, but no known biosynthetic products (Pearson et al., 2007). To reconcile such orphan biomarkers and mystery cyclases, we analyzed the modern diversity and evolutionary history of the terpenoid cyclase protein superfamily, of which triterpenoid cyclases are one protein family.

**Figure 1:**
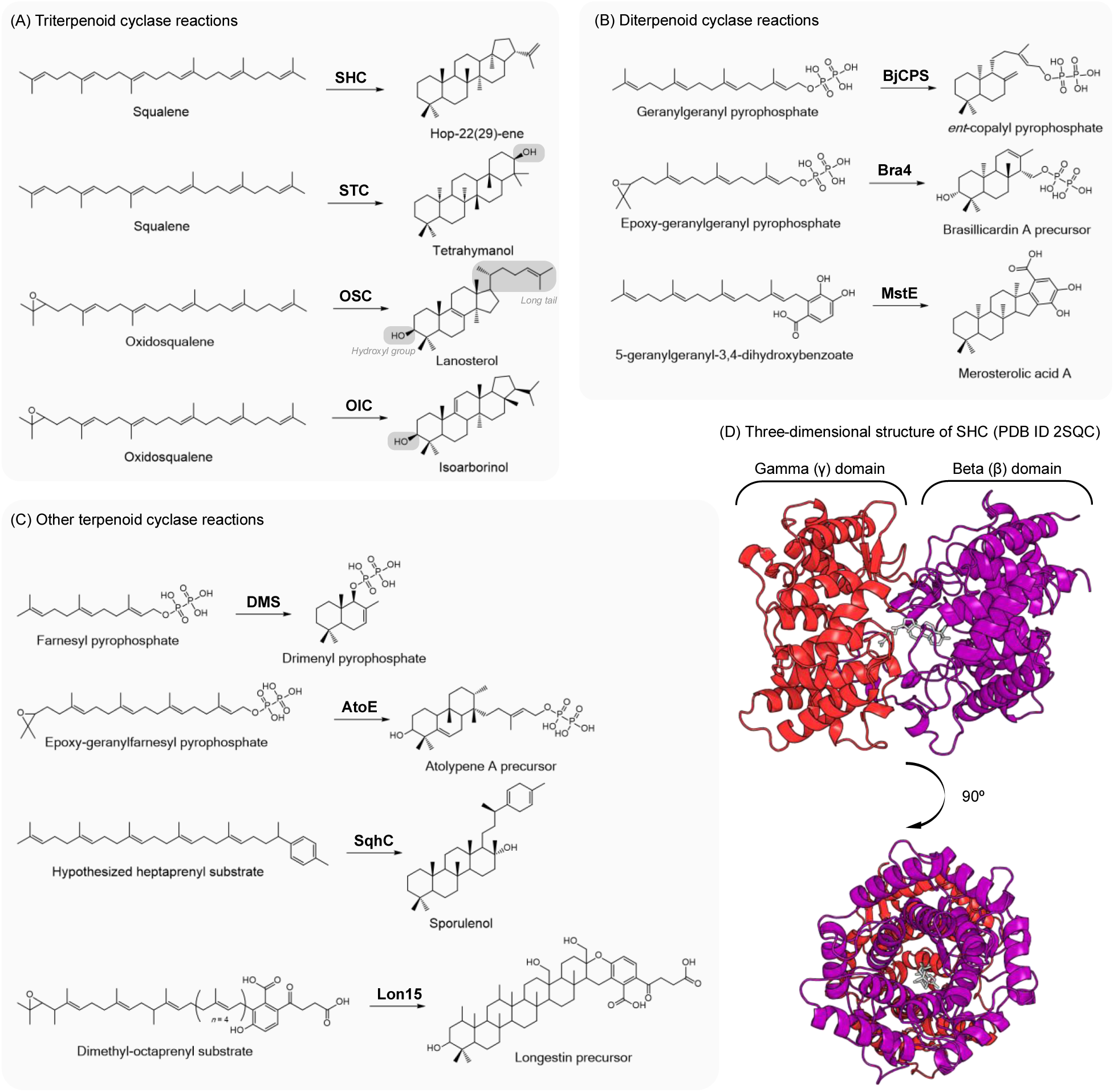
Reactions performed by Type II terpenoid cyclases, including on (A) C_30_ substrates in the case of triterpenoid cyclases (with annotation showing important chemical differences among products), (B) C_20_ substrates in the case of diterpenoid cyclases, and (C) C_15_, C_25_, C_35_, and C_40_ substrates for other Type II terpenoid cyclases. Other reactions exist for some substrates; for example, in addition to lanosterol cyclases, there are oxidosqualene cyclases that synthesize parkeol, cycloartenol, or amyrins. (D) The crystal structure of squalene-hopene cyclase from *Alicyclobacillus acidocaldarius* (Wendt et al., 1999). SHC: squalene-hopene cyclase; STC: tetrahymanol cyclase; OSC: oxidosqualene cyclase; OIC: oxidosqualene-isoarborinol cyclase; BjCPS: *Bradyrhizobium japonicum* copalyl diphosphate synthase; Bra4: brasilicardin cyclase; MstE: merosterolic acid synthase; DMS: drimenol synthase; AtoE: atolypene synthase; SqhC: sporulenol synthase; Lon15: terpene cyclase (longestin).

Terpenoid cyclases rearrange carbon-carbon bonds to form polycyclic molecules through an exquisitely precise carbocation cascade (Abe et al., 2014; Eschenmoser & Arigoni, 2005;Hoshino & Sato, 2002). We refer here specifically to the Type II terpenoid cyclase protein family, which protonates substrates with a general acid, and is distinct from the Type I terpenoid cyclase family, which adopts a different fold and employs a different, metal-dependent mechanism to cyclize C_10_, C_15_, and C_20_ substrates (Christianson, 2017). Type II terpenoid cyclases typically have a dumbbell shape composed of two homologous structural domains known as the β and γ domains. The enzyme active site is between the two domains, accessible by a substrate channel through the γ domain, while the β domain has a catalytic aspartic acid, and a water tunnel needed to replenish it. Some Type II cyclases have additional domains or lack the γ domain. All domains share the alpha-alpha toroid fold, which is a torus of 12 alpha helices arranged in two concentric rings of 6 (Figure 1). As described by the Structural Classification of Proteins – extended 2.08 (Chandonia et al., 2022; Fox et al., 2014), terpenoid cyclases and protein prenyltransferases make up one of the six superfamilies in the alpha-alpha toroid fold, the others being glycosidases and polysaccharide lyases (Syrén et al., 2016), an enzyme involved in the synthesis of lanthipeptide antibiotics (Zhang et al., 2012), and eukaryotic α2-macroglobulins and complement proteins (Cao et al., 2010).

Type II terpenoid cyclases include triterpenoid (C_30_ substrate) and diterpenoid (C_20_ substrate) cyclases, which share a catalytic mechanism and βγ domain architecture (Christianson, 2017). Triterpenoid cyclases include oxidosqualene cyclases, which cyclize an epoxidized substrate to form tetracyclic sterols such as lanosterol, parkeol, cycloartenol, arborinols, and amyrins (R. Xu et al., 2004). These lipid alcohols are often modified to form functional membrane components such as cholesterol. Squalene-hopene cyclases and tetrahymanol cyclases cyclize squalene to form pentacyclic hopenes and tetrahymanol, respectively. Unmodified tetrahymanol functions as a membrane lipid (Conner et al., 1971), and hopanoids most often undergo polyfunctionalization at the “tail” alkene to become amphipathic membrane components as well (Belin et al., 2018; Bradley et al., 2010). Diterpenoid cyclases cyclize geranylgeranyl pyrophosphate (C_20_) to form bicyclic copalyl (Ikeda et al., 2007; Morrone et al., 2009; Smanski et al., 2011), halimadienyl (Nakano et al., 2005), terpentedienyl (Dairi et al., 2001; Stowell et al., 2022), and kolavenyl (Nakano et al., 2015) diphosphates, among other molecules. Other diterpenoid cyclases work on epoxidized geranylgeranyl pyrophosphate, forming tricyclic molecules such as brasilicardin (Hayashi et al., 2008), phenalinolactone (Dürr et al., 2006), and tiancilactone (Dong et al., 2018) precursors. As shown in Figure 1, the protein superfamily includes the above and also C_15_ (Pan et al., 2022; Vo et al., 2022), C_25_ (Kim et al., 2019), C_35_ (Kontnik et al., 2008; Sato, Hoshino, et al., 2011; Sato, Yoshida, et al., 2011), and C_40_ (Hayashi et al., 2007; Ozaki et al., 2018) cyclases, which work on a variety of alkenyl, epoxidized, and otherwise functionalized isoprenoid substrates.

Products of triterpenoid cyclases often undergo modification by other enzymes and go on to regulate membrane homeostasis and dynamics, among other functions (Bloch, 1983). The best-studied example is cholesterol in the animal cell membrane, where it buffers membrane flexibility against temperature changes (Crockett, 1998; Mouritsen & Zuckermann, 2004) and organizes the membrane into liquid-ordered microdomains (Levental et al., 2020; Xu & London, 2000). There is evidence for similar function for hopanoids in bacteria (López & Kolter, 2010; Sáenz, 2010; Sáenz et al., 2012, 2015). Cyclic diterpenoids, on the other hand, are commonly modified with solubilizing functional groups after cyclization, and function in cell-cell and interspecific communication. Meanwhile, diterpenoids cyclized by Type I cyclasesare often insoluble and are involved in other functions such as plant wound healing in the form of resins. Other well-studied diterpenoids include the gibberellin hormones made by plants and their symbionts to regulate tissue growth (Nett et al., 2022). Cyclic diterpenoids are also involved in interspecies conflict – between microbes in the form of antibiotics such as terpentecin (Tamamura et al., 1985), platencin (Jayasuriya et al., 2007; J. Wang et al., 2007), platensimycin (J. Wang et al., 2006), and phenalinolactone (Gebhardt et al., 2011), antifungals such as viguiepinol and oxaloterpins (Bi & Yu, 2016), and between humans and their pathogens in the case of tuberculosinol from *Mycobacterium tuberculosis* (Mann et al., 2009) and brasilicardin A from *Nocardia brasiliensis* (Shigemori et al., 1998; Usui et al., 2006).

The evolutionary processes that generated the structural and functional diversity of terpenoids and of triterpenoids in particular have long been a subject of great scientific interest (Bloch, 1983; Ourisson, 1989; Ourisson & Nakatani, 1994). Fischer and Pearson (2007) presented several alternative models for relationships among triterpenoid cyclases under parsimonious evolution of three functional traits, which they posit evolve independently and slowly, if not irreversibly: substrate (squalene vs. oxidosqualene), reaction favorability (number of anti-Markovnikov carbocations propagated during the reaction), and product stereochemistry (whether the first three rings are chair-chair-chair vs. chair-boat-chair). In this study, we adopt their rigorous evolutionary approach and update it with four new sources of information, in pursuit of the same biological question: in what order, and through what processes, did the diversity of polycyclic membrane regulators arise?

We first apply a probabilistic method (maximum likelihood) of estimating relationships among cyclases, using an explicit model of protein sequence evolution. This allows us to relax Fischer and Pearson’s (2007) assumptions about the independence of the traits and the (slow) speed at which they evolve. Therefore, we can observe the distribution of the selected cyclase traits across a tree generated under unrelated assumptions (amino acid exchangeabilities and rate distributions) rather than using these trait assumptions to build the tree. We also revise the list of traits based on functional studies of cyclases over the past 16 years. Our use of probabilistic phylogenetic methods follows that of previous workers (Desmond & Gribaldo, 2010; Frickey & Kannenberg, 2009; Gold et al., 2017; Santana-Molina et al., 2020), and we find that our analyses, where they overlap, recapitulate theirs. Second, we analyze the astounding cyclase diversity since sequenced, with special attention to divergent sequences from environmental metagenomes and metagenome-assembled genomes as a potential source of cyclases that synthesize orphan biomarkers or novel natural products. Previous environmental sequencing efforts focused on triterpenoid cyclases found entire clades comprised solely of metagenomic sequences (Pearson et al., 2007; Pearson & Rusch, 2009), all of which remain uncharacterized. Third, we broaden the analysis to include the entire terpenoid cyclase protein superfamily allowing us to root the triterpenoid cyclase protein family tree, again independently of assumptions about the manner in which functional traits evolve. Fourth and finally, using the evolutionary framework provided by an explicit model of protein evolution, sequence diversity, and the alpha-alpha toroid fold, we identify cyclases that branch at key transitions in the superfamily’s evolution. We test the function of these cyclases via heterologous expression in *Escherichia coli* and find two surprising enzymes. One is a cyclase that synthesizes isoarborinol, an orphan biomarker from the Paleozoic. The other is a group of cyclases within the squalene-hopene cyclase clade which cyclize oxidosqualene (the native sterol cyclase substrate) but not squalene (the native squalene-hopene cyclase substrate). In addition to other lines of evidence, the substrate specificity of these cyclases suggests that alkenyl substrates may not be ancestral to triterpenoid cyclases, and were gained relatively late in the squalene-hopene cyclase lineage.

## Results and Discussion

### Phylogeny

We began our analyses of cyclase evolution by estimating the phylogeny of all proteins known to share the cyclase fold, including triterpenoid, diterpenoid, and meroterpenoid cyclases, and cyclase homologs of unknown function from environmental metagenomes (Figure 2). To do so, we retrieved cyclase homologs from public databases using BLAST and Pfam-based methods (see Material and Methods). Combined, these searches resulted in 44,209 unique sequences after removing duplicates and filtering for length. Briefly, all sequences in this database were aligned, alignments were subset to 1,139 sequences, and phylogenetic trees were estimated under maximum likelihood.

**Figure 2:**
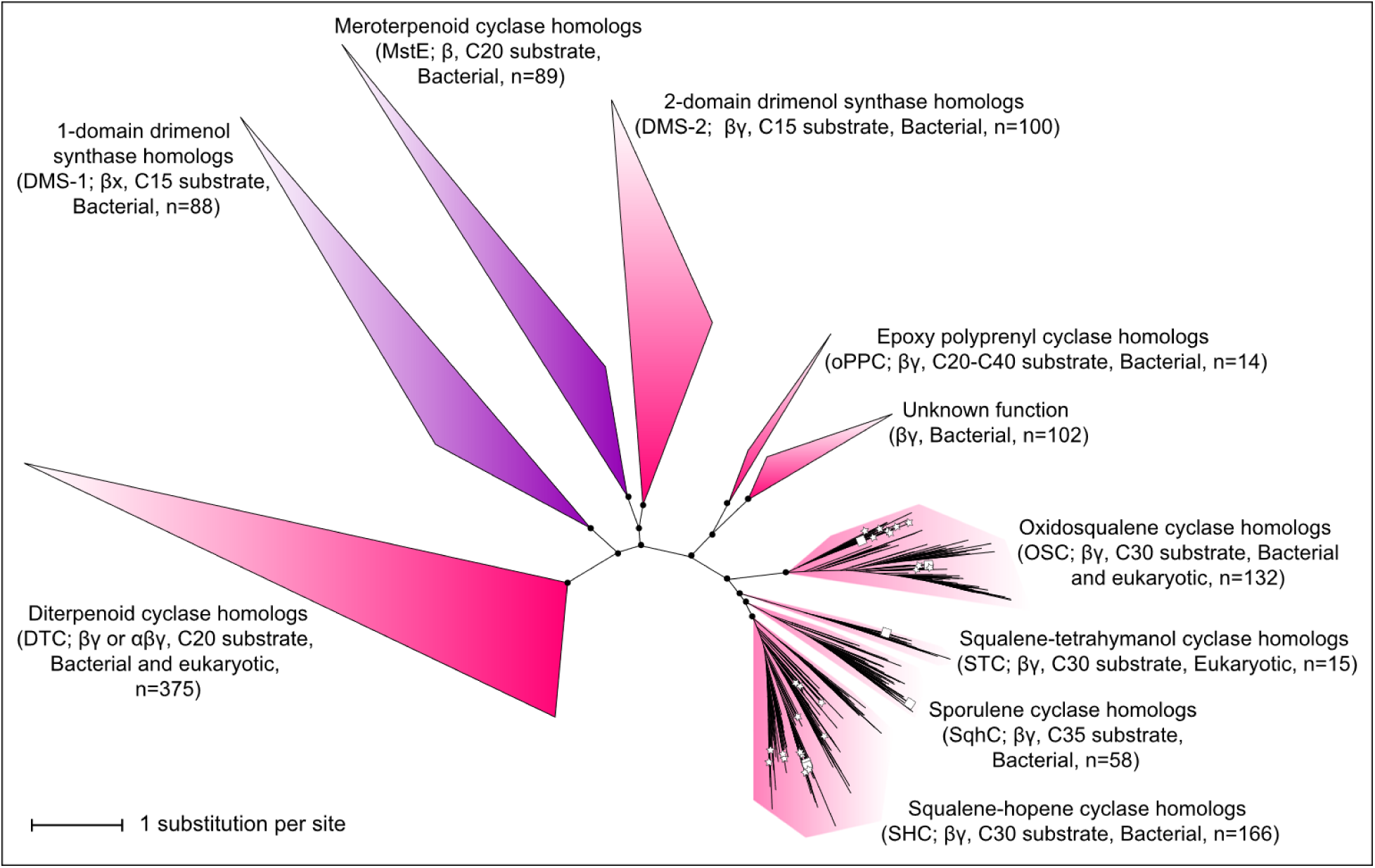
Maximum-likelihood phylogeny of terpenoid cyclases estimated under the EX_EHO model with 5 free rate categories. Stars indicate cyclases from environmental metagenomes; squares indicate cyclases of known function. Ultrafast bootstrap support is shown for deep splits, where black circles indicate bootstrap ≥ 97. Purple clades are composed of proteins without a γ domain (β-only or βx) as annotated; pink clades are composed of two-domain (βγ or αβγ) proteins.

We tested the robustness of tree topology to uncertainty in alignment, model choice, and history of domain duplication and loss. Figure 2 shows the cyclase phylogeny estimated under EX_EHO+R5, which applies separate evolutionary models to each site depending on its probability of being on the protein’s surface or buried, and depending on whether it is in an alpha helix, a loop, or another structural element (Le & Gascuel, 2010). This model also calculates the probability of each site being in one of 5 evolutionary rate categories. While the distribution of these categories differed slightly between the β and γ domains, we found that the likelihood-maximizing partitioning scheme for the concatenation of domains was a single partition. The topology estimated under EX_EHO+R5 is robust to omission of γ domains, to alignment masking, to larger and smaller subsets of the sequence database, and to model misspecification, as the same deep splits were recovered under runner-up models (Supplementary Table 2). However, ultrafast bootstrap support improved dramatically under better-fitting models. Thus, although protein sequences contain limited evolutionary information, and can lose information as they evolve due to site saturation caused by repeated substitution (Xia et al., 2003), the phylogeny presented here represents a strong hypothesis given the sequence information available.

The domain structure of all cyclases in this tree is either β, βγ, βx, or αβγ. We noticed that β and γ domains crystal structures do not align well, despite clear secondary structural and topological homology (Supplementary Table 1). Similarly, diterpenoid γ domains and triterpenoid γ domains do not align well, due either to divergence or because they arose from independent duplications of the β domain. If so, the diterpenoid and triterpenoid cyclase γ domains would still be homologous but could have arisen from β domains with different sequences, in different organisms, resulting in different trajectories through fitness landscapes and resultingly independent evolutionary constraints, rates, and substitution processes. For this reason, we used structural alignments of protein crystals to create custom hidden Markov models (HMMs) for the β domain, the diterpenoid cyclase γ domain, and the triterpenoid cyclase γ domain. Our triterpenoid γ domain HMM did not pick up the diterpenoid γ domain, and vice versa. We estimated cyclase phylogeny with the two γ domains aligned separately and recovered the same topology as when γ domains were aligned together, or omitted.

### Major clades

The overall tree topology is consistent with previously published triterpenoid and diterpenoid subtrees (Desmond & Gribaldo, 2010; Frickey & Kannenberg, 2009; Gold et al., 2017; Santana-Molina et al., 2020), and with a recent tree of all terpenoid cyclases (Hoshino & Villanueva, 2023). Squalene-hopene cyclase homologs (SHCs) and oxidosqualene cyclase homologs (OSCs) form large monophyletic clades, with squalene-tetrahymanol cyclase homologs (STCs) from microbial eukaryotes and sporulenol synthase homologs (SqhCs) from Bacillota forming small groups sister to SHC, and bacterial OSC groups branching earliest among the OSC subclades.

Sister to this mostly-triterpenoid group is another pair of clades, one of which is a small group of epoxy-polyprenyl cyclases. These enzymes take a substrate that is epoxidized like oxidosqualene but has all head-tail joined isoprene units rather than the central tail-tail linkage of squalene and oxidosqualene (Figure 1). The epoxy-polyprenyl cyclases include Lon15, a meroterpenoid octaprenyl (C_40_) cyclase from Actinomycete *Streptomyces argenteolus* (Hayashi et al., 2007) and AtoE, a unique C_25_ cyclase from Actinomycete *Amycolatopsis tolypomycina* NRRL B-24205 (Kim et al., 2019). This clade is quite taxonomically restricted, with most sequences coming from Actinomycetes and a handful from Chloroflexota and Clostridia. Surprisingly, those diterpenoid cyclases that utilize an epoxidized substrate – Bra4 and PlaT2, epoxy-diterpenoid cyclases from Actinomycetes *Nocardia brasiliensis* IFM 0406 (Hayashi et al., 2008) and *Streptomyces* sp. Tü 6071 (Dürr et al., 2006), respectively – also fall in this clade rather than with other diterpenoid cyclases. It seems that the number of substrate isoprenoid chain carbons is a polyphyletic trait in the cyclase tree, as demonstrated by these diterpenoid cyclases and MstE (which takes a substrate with a dihydroxybenzoate-functionalized C20 isoprenoid chain) and by the triterpenoid cyclases, which are paraphyletic with respect to C35 SqhC. The position of SqhC also demonstrates the relative lability of isoprenoid chain linkage (all head-tail vs. tail-tail) as a trait.

Sister to the epoxy-polyprenyl cyclase group is a clade of unknown function referred to in the literature (e.g., Santana-Molina et al., 2020) as “SHC-like.” This clade contains Gracilicutes sequences from the PVC group (Planctomycetes, Verrucomicrobia, Chlamydiae), the FCB group (Fibrobacterota, Chlorobiota, Bacteroidota), Spirochaetota, and Acidobacteriota. The function of these proteins, including those from cultured and genetically tractable organisms such as *Gemmata obscuriglobus,* has not been demonstrated, and their molecular structure has not been determined. However, we found that for both groups, the γ domains are detectable by the custom “triterpenoid” γ domain HMM we generated but not by our “diterpenoid” γ domain HMM. This could be due to functional divergence of the γ domains in these two halves of the tree (see “evolutionary implications” below).

On the tree’s central long branch are three subgroups, each containing a single characterized enzyme. One is composed of monodomain (β-only) meroterpenoid cyclase homologs, including a single characterized enzyme from Cyanobacterium *Scytonema sp.* PCC 10023, which cyclizes a quinone precursor with a functionalized benzyl headgroup and a C_20_ isoprenoid chain (Moosman et al. 2020). Interestingly, this cyclase, called MstE, uses a DxD motif rather than the canonical DxDD motif to protonate an alkenyl substrate. Other sequences in this clade are from diverse bacterial taxa, including other Cyanobacteria as well as Alphaproteobacteria, Betaproteobacteria, Deltaproteobacteria, Gammaproteobacteria, Acidobacteriota, Myxococcota, Planctomycetes, Bacillota, Bacteroidota, Actinomycetota, and Chloroflexi, spanning the Gracilicutes and Terrabacteria divisions, although not recapitulating their branching order. There are also several archaeal sequences, including from cultured representatives such as *Methanosarcina acetivorans* C2A. The other subgroups are composed of drimenyl pyrophosphate synthases (DMS), recently-discovered enzymes that cyclize a C15 substrate. One (“DMS-2”) contains the two-domain DMS characterized from Actinomycete *Streptomyces showdoensis* (Pan et al., 2022), along with homologs from other bacteria, while the other (“DMS-1”) contains the DMS characterized from Bacteroidete *Aquimarina spongiae* (Vo et al., 2022), which has only one alpha-alpha toroid domain, fused to a haloacid dehalogenase domain. This clade also contains homologs from Actinomycetota, FCB group bacteria, and fungi.

The final major clade consists of diterpenoid cyclases. All characterized proteins in this group take geranylgeranyl pyrophosphate (GGPP) as a substrate. The two most basal clades consist of only bacterial proteins, including characterized bacterial cyclases that make *ent*-copalyl pyrophosphate (Jayasuriya et al., 2007; Morrone et al., 2009; Smanski et al., 2011), a gibberellin or antibiotic precursor, and the cyclase from *Kitasatospora griseola* that synthesizes the terpentecin precursor (Dairi et al., 2001). This is sister to a pair of clades containing fungal *ent*-copalyl pyrophosphate cyclases (Quin et al., 2014) and plant *ent*-copalyl pyrophosphate cyclases (Jia et al., 2022), respectively. The plant clade has many paraphyletic groups of prokaryotic homologs on its stem, including the *Mycobacterium tuberculosis* cyclase that synthesizes the tuberculosinyl adenosine precursor (Young et al., 2015) and homologs from the Asgardarchaeota that produce the same molecule (McShea et al., 2025).

### Functional analysis of putative metagenomic cyclases

Cyclases from environmental metagenomes are distributed throughout the tree, especially in bacteria-dominated clades (Figure 3). They fall in the SHC and OSC crown groups, as well as on the stems of these clades, in small groups or as singleton branches. To determine if these metagenomic cyclases are capable of cyclizing squalene or oxidosqualene, we expressed a select set of cyclases in two strains of *E. coli,* engineered to synthesize squalene or oxidosqualene, respectively, and measured production of polycyclic lipids by gas chromatography-mass spectrometry (GC-MS). We identified 14 triterpenoid cyclase homologs from environmental metagenomes which either branched on the OSC or SHC stem lineage rather than within the crown group or branched within the crown in monophyletic subgroups with no cultured representatives. Many of these cyclases have substitutions at sites of known importance for cyclization of squalene or oxidosqualene.

**Figure 3:**
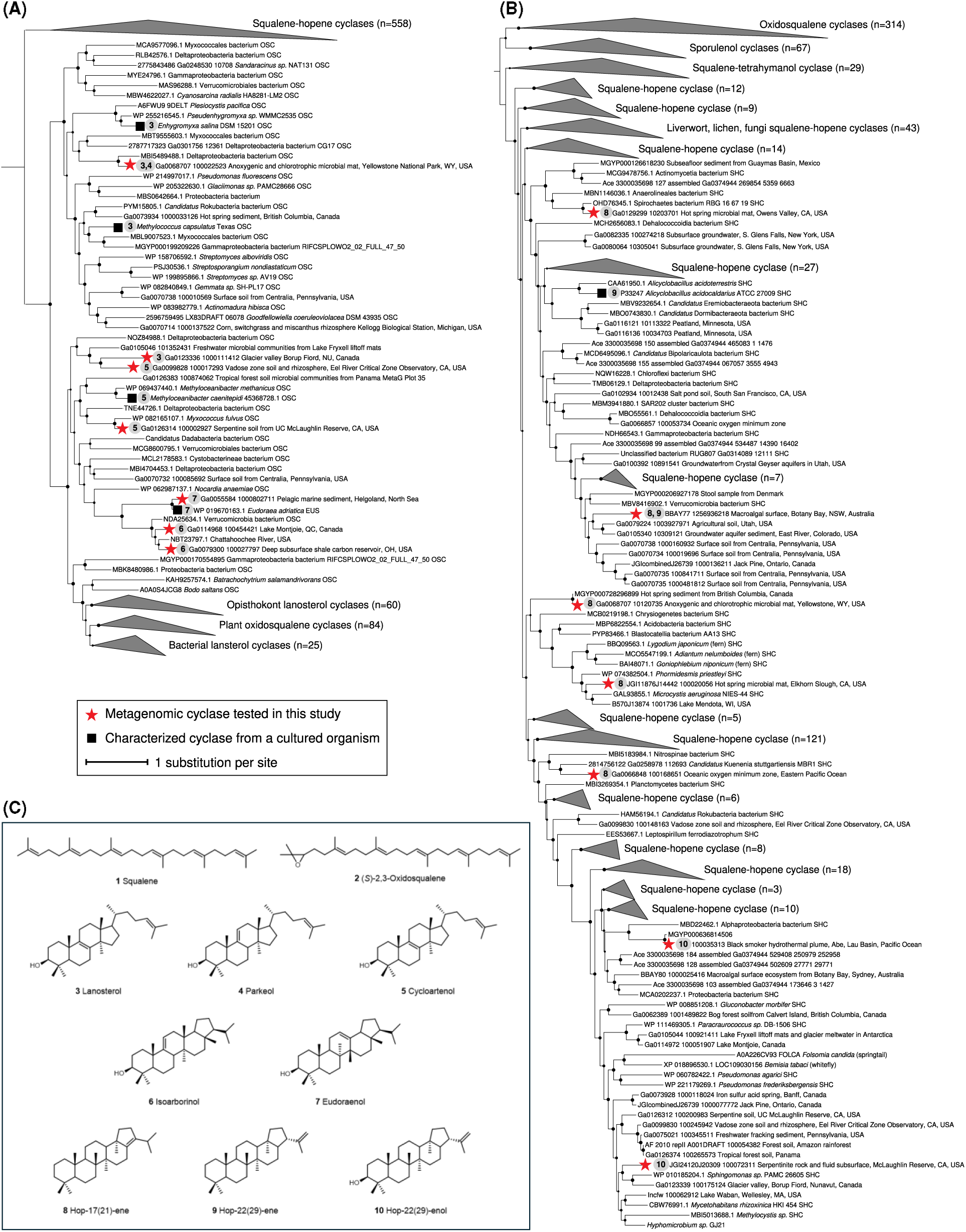
Phylogenetic positions of triterpenoid cyclases from environmental metagenomes heterologously expressed in *E. coli*. Metagenomic cyclases are marked with red stars, while previously characterized enzymes from cultured organisms are marked with black squares, and both are annotated with product profiles. (A) Oxidosqualene cyclases. (B) Squalene-hopene cyclases. (C) Substrates and products of cyclases from environmental metagenomes.

Many triterpenoid cyclases were active in the heterologous expression system. Most enzymes from early-branching OSC subclades produce lanosterol or cycloartenol (Table 1; structures shown in Figure 3C), consistent with their phylogenetic proximity to cyclases from cultured bacteria known to make those products (Figure 3A). None of the cyclases from early-branching bacterial OSC groups had activity for squalene, which requires the greater active site acidity of an SHC (Xu et al., 2004).

**Table 1:**
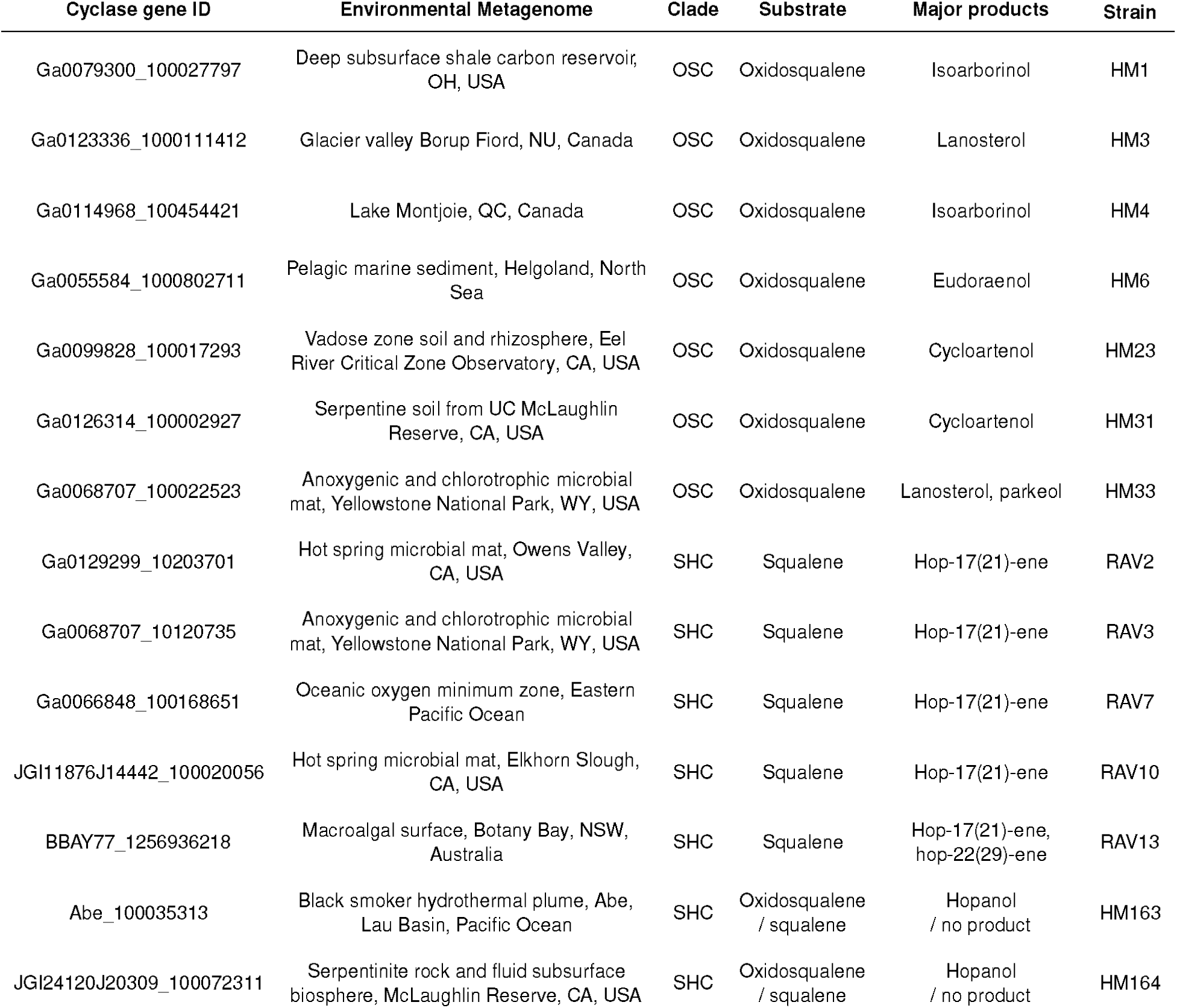
Substrates and products of triterpenoid cyclases from environmental metagenomes heterologously expressed in *E. coli*. Products were determined by comparison to published GC-MS spectra (see Supplementary Figures 1 and 2), and in the case of isoarborinol, confirmed by NMR (Supplementary Figure 3 and Supplementary Table 3).

One cyclase from a marine metagenome produces eudoraenol, a recently discovered arborane triterpenoid. It is the second eudoraenol-synthesizing enzyme to be reported after the discovery of a eudoraenol cyclase in *Eudoraea adriatica* (Banta et al., 2017), a marine heterotroph from phylum Bacteroidetes (Alain et al., 2008). The new eudoraenol cyclase has the same substitutions shown by Banta et al. (2017) to be sufficient for eudoraenol production – tryptophan to serine at position 230 (*Homo sapiens* OSC numbering), histidine to tyrosine at position 232, tyrosine to valine at position 521, and asparagine to tyrosine at position 697. In the enzyme active site, these residues are clustered near the substrate tail where the fifth ring forms in arborinols.

Closely related to the eudoraenol cyclases are two cyclases from deep subsurface and inland lake ecosystems which we found synthesize isoarborinol (Figure 4). These cyclases have the eudoraenol cyclase substitutions at positions 230, 232, and 697 and a unique cysteine at position 503. The subsurface cyclase contig is predicted by the Whokaryote classifier (Pronk & Medema, 2022) to be prokaryotic, making it the first reported bacterial source of isoarborinol. While the lake cyclase contig is too short for classification, its phylogenetic proximity to the subsurface and other bacterial cyclases make it likely to be prokaryotic as well. Arborinol is named for the tree from which it was first isolated (Kennard et al., 1965; Vorbrüggen et al., 1963), but the arborane fossil record stretches back to the Permian (Hauke et al., 1995), well before the evolution of angiosperms, leading to the hypothesis (Ourisson et al., 1982) that microbial sources of arborinols, and specifically isoarborinol, must exist. The formation of isoarborinol by these two cyclases identifies a modern and most likely bacterial source for this “orphan” biomarker. Bacterial arborinol cyclases form a monophyletic clade within one of the bacterial OSC clades. Thus, it is unclear whether bacterial arborinol cylcases arose before or after the radiation of eukaryotic OSCs, which show a pattern of vertical inheritance (i.e., major clades of plant, fungal, and animal cyclases largely recapitulate organismal phylogeny). The bacterial arborinol cyclases are thus a reasonable, but not definitive, source for Paleozoic and Proterozoic arborane fossils.

**Figure 4:**
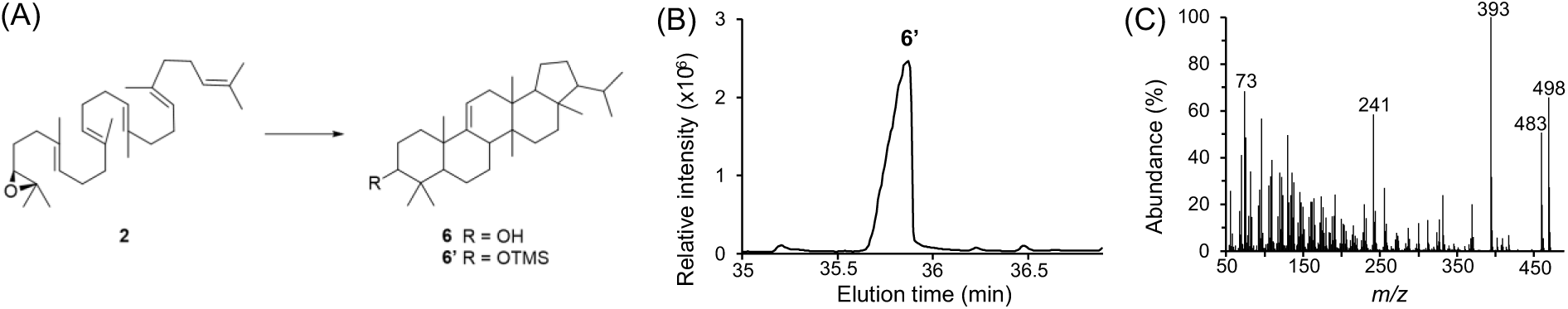
Biosynthesis of isoarborinol by a triterpenoid cyclase from a deep subsurface shale metagenome. (A) The reaction performed by isoarborinol cyclase. (B) GC-MS total ion chromatogram of total lipid extract from *E. coli* strain HM1 expressing the oxidosqualene biosynthesis pathway and Ga0079300_100027797, derivatized to trimethylsilyl (TMS) ethers. (C) Mass spectrum of peak at 36.0 min, identified by comparison to Vorbrüggen et al. (1963) and by nuclear magnetic resonance (NMR), see Supplementary Figure 3 and Supplementary Table 3.

Most enzymes from early branching SHC subclades (Figure 3B) that we tested produce typical hopene isomers hop-17(21)-ene and/or hop-22(29)-ene from a squalene substrate (Table 1). Hopene-producing enzymes also all had activity for oxidosqualene, typically producing a single hopanol isomer (hopan-22(29)-ol), consistent with previous work which has shown squalene cyclases to be nonspecific (Abe & Rohmer, 1994; Hammer et al., 2013; Rohmer et al., 1980; Seitz et al., 2012). However, a small group of cyclases within the SHC crown had activity for only oxidosqualene and no activity for squalene, producing only hopanols. This is surprising given that all other cyclases in the SHC clade have some activity for squalene, including sporulenol cyclases, which are functional for both β-curcumene and squalene, and tetrahymanol cyclases. These oxidosqualene cyclases in the SHC clade do not lack any residues needed to activate squalene. Indeed, they have the electrophilic DxD[D] motif thought to be necessary to protonate an alkenyl substrate. It is possible that these enzymes are in fact true SHCs but cannot cyclize squalene in the *E. coli* cell under the conditions tested, perhaps due to minor protein misfolding that weakens the electrophilicity of the active site, retaining activity for the more reactive substrate but abolishing it for the more recalcitrant one. It is also possible that these hopanol-producing enzymes are true OSCs in the functional sense, despite being SHCs in the phylogenetic sense, and that substrate functionalization is a relatively labile trait in this group.

### Adaptation in the terpenoid cyclase superfamily

What evolutionary dynamics drove the diversification of terpenoid cyclases? Did the diversity of substrates and products in this superfamily arise from neutral divergence and drift, or did natural selection play a role? To assess the relative influences of drift and selection, we employed a quantitative test that interprets positive selection as an elevated rate of amino acid mutations in functional regions of the protein structure, along a given branch in a phylogenetic tree (similar to Maddamsetti & Grant, 2022; Ritchie et al., 2021). This approach applies the theory developed for protein-coding nucleotide sequences (e.g. the branch-site test, Gharib & Robinson-Rechavi, 2013; McDonald & Kreitman, 1991; Messer & Petrov, 2012; Yang & Dos Reis, 2011) to protein structures, which allows the test to work on deeply divergent groups where nucleotide substitutions have saturated but structure has evolved more slowly.

Along all the major internal branches of the terpenoid cyclase phylogeny, there are two where a functional domain had a significantly elevated proportion of amino acid substitutions over background (Figure 5). One is the stem of the triterpenoid cyclases which take an alkenyl substrate (that is, squalene-hopene, sporulenol, and tetrahymanol cyclases), where the active site of the protein has an unusually high substitution density. The second is the stem of the group of unknown function, where the SHC-type dimer contact surface had unusually high substitution density. We interpret clustering of substitutions in a functional region as a potential signal of protein adaptation, in which case these may be two instances of positive selective pressure and/or functional divergence in the evolutionary history of triterpenoid cyclase homologs. Adaptation could be to new environmental conditions (e.g. to a new host organism after horizontal gene transfer), to new function, or both. That only two instances of rate shifts in functional regions were detectable means either that adaptation on other branches occurred at a molecular length scale undetectable when focusing on functional regions (e.g. single-residue substitutions), at a temporal scale undetectable when focusing on long branches (e.g. rapid evolution within crown groups), or that triterpenoid cyclase divergence was otherwise dominated by drift.

**Figure 5:**
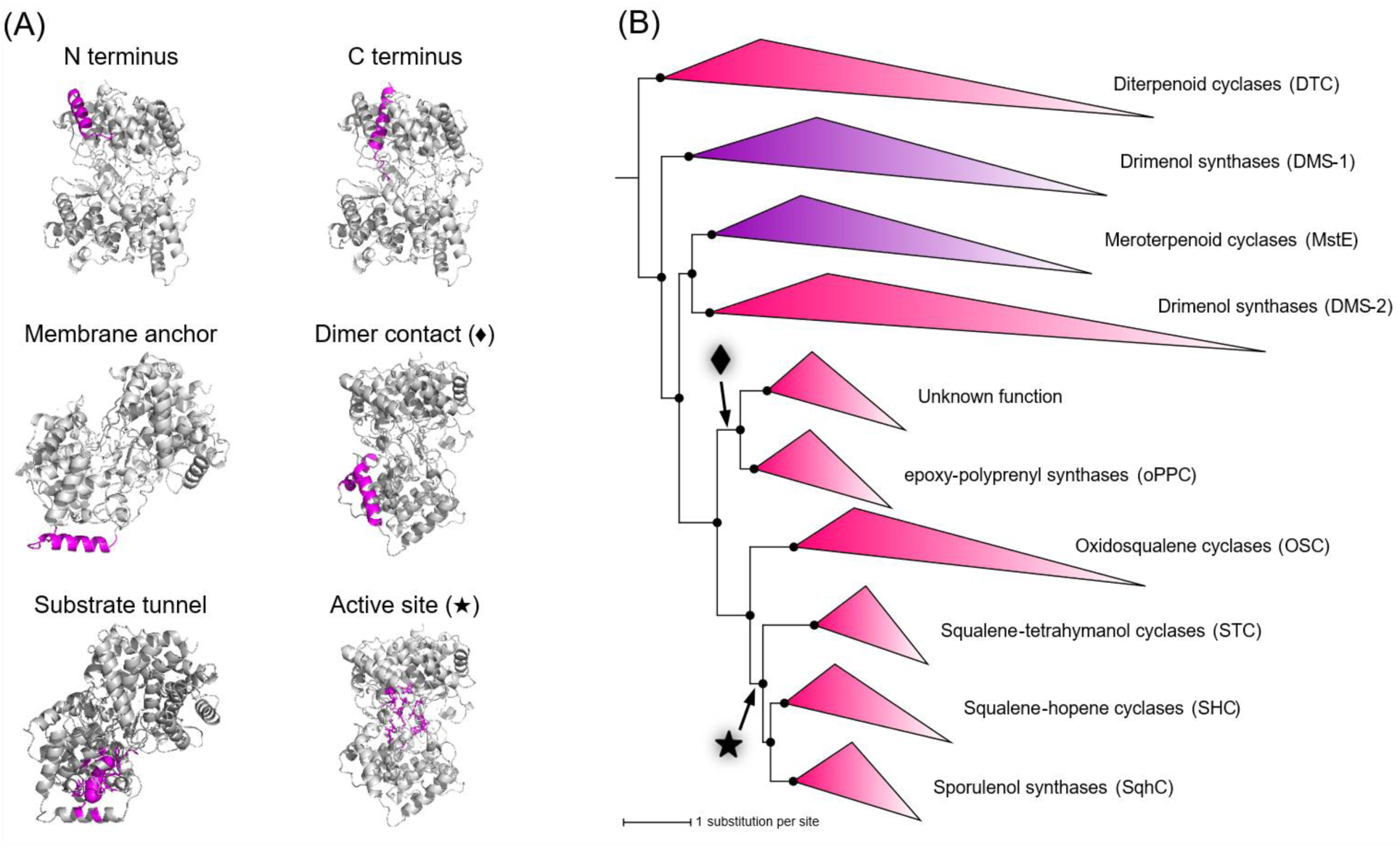
Windows of rapid evolution in functionally important domains of triterpenoid cyclases. (A) Functional regions shown on the squalene-hopene cyclase structure (PDB ID 1SQC). Regions found to have an evolutionary rate elevated above background are marked with a rhombus (⧫) and a star (★). (B) Branches with a rapidly-evolving functional region, marked with symbols corresponding to panel A. As in Figure 2, purple clades are composed of proteins without a G domain (β-only or βX); pink clades are composed of two-domain (βγ or βγX) proteins.

### Evolutionary Implications

sTo interrogate domain evolution in the cyclase family, we turned to protein structure, which evolves more slowly than sequence composition (Caetano-Anollés et al., 2009; Ingles-Prieto et al., 2013). We applied structural data to estimate relationships in three ways: we used structures to better align sequences before estimating phylogeny under a model of sequence evolution (Figure 6A), we measured relationships as simple distances between structural coordinates (Figure 6B), and we measured relationships using the local structural alphabet implemented in Foldtree (Moi et al., 2023) (Figure 6C).

**Figure 6:**
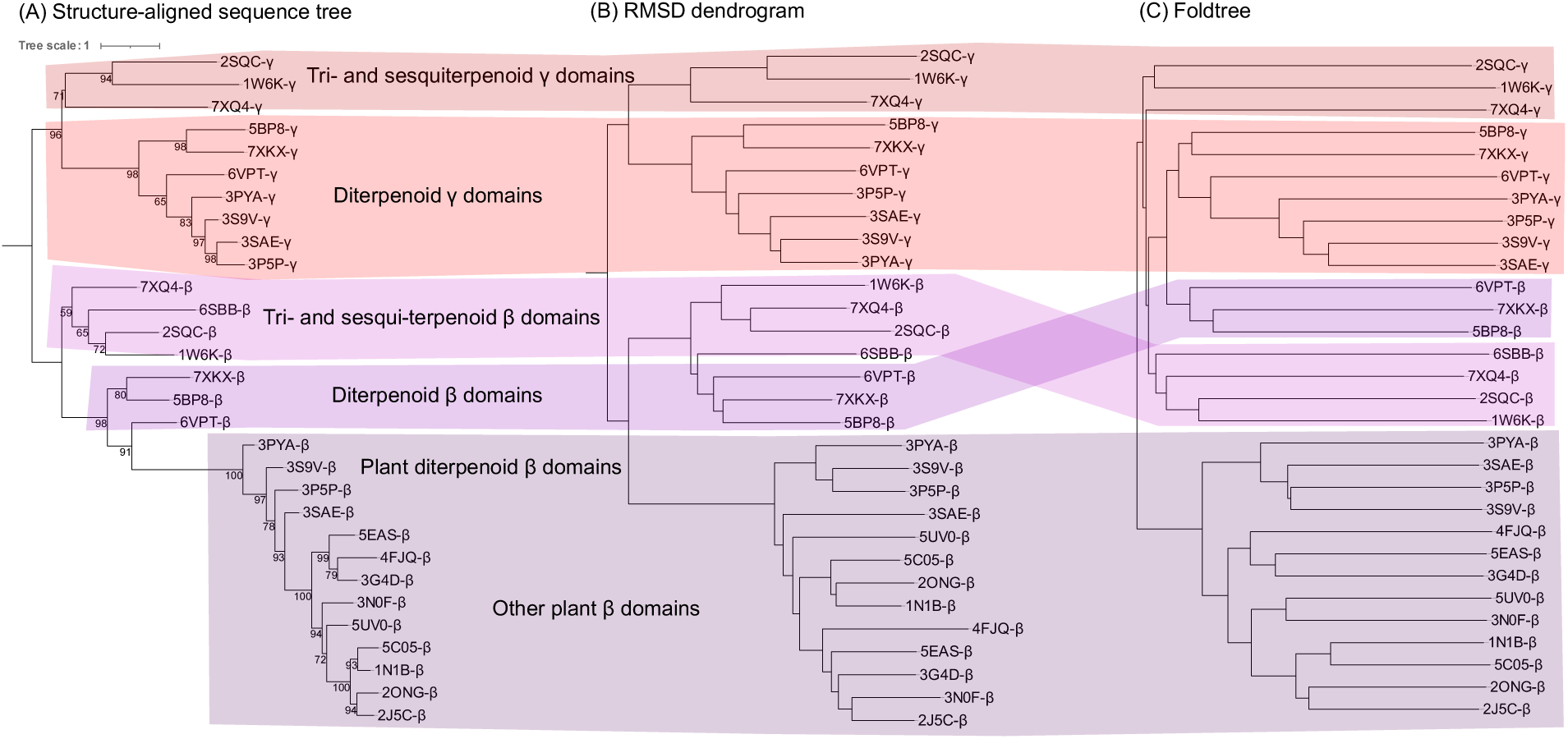
Relationships between cyclase domains estimated using three structure-informed methods. (A) Phylogeny estimated using standard sequence-based method after aligning sequences using structure. (B) Neighbor-joined dendrogram of domains using root mean squared deviation (RMSD) as distances. (C) Structure relationships estimated using the Foldtree algorithm. Red and purple stripes are for visual aid, to show the position of each group across trees.

The structure-aligned sequence tree (Figure 6A) and the RMSD dendrogram (Figure 6B) show a deep split between β domains and γ domains. The β and γ clades have the same internal branching order in the structure-aligned sequence tree, consistent with a single duplication event at the root followed by parallel divergence, with the only other domain rearrangements being the loss of the γ domain in monodomain cyclase MstE (PDB ID 6SBB), gain of the α domain in the ancestor of plant cyclases, and loss of the γ domain later within that clade. The RMSD dendrogram also supports this scenario, and differs only in the relationships among β domains, placing the β domains of bacterial diterpenoid cyclases sister to those of the triterpenoid cyclases rather than those of the plant cyclases. The Foldtree estimate (Figure 6C) predicts a much more complex scenario, with a deep split between plant β domains and the rest of the structures. After the split with plant βs, the first branches are triterpenoid and sesquiterpenoid γ domains, then tri- and sesqui-terpenoid βs, then diterpenoid βs and γs. This is consistent with our hypothesis of triterpenoid and diterpenoid β domains arising from independent duplication events but suggests a surprising γ domain ancestor for all non-plant β domains. Given that Foldtree does not yet measure statistical support, and given the agreement between the structure-aligned sequence tree, the RMSD dendrogram, and our large sequence-based phylogeny (Figure 2), we tentatively reject the Foldtree estimate and the independent duplications hypothesis. Most evidence supports a single evolution of the terpenoid cyclase γ domain. However, this scenario should be revisited as structural phylogenetics (Garg & Hochberg, 2025; Malik et al., 2020; Moi et al., 2023) becomes more sophisticated (Mutti et al., 2025).

Given the possible selection on oligomerization and the active site revealed through our structural analyses of cyclases, we can posit and assess evolutionary scenarios focused on related traits. The longest internal branch of the cyclase phylogeny is between the diterpenoid cyclases and all other proteins (Figure 7A). Rooting the tree on this branch, in agreement with the root position in the β and γ domain subtrees (Figure 6), places all-bacterial clades basal to the eukaryotic clades. This is consistent with separate transfer events of diterpenoid cyclases from bacteria to plants and to fungi, likely at different times and involving different donor organisms and proteins. For this and most possible superfamily roots, the triterpenoid cyclase root is between the squalene-hopene (and STC and SqhC) and oxidosqualene cyclases. This OSC-SHC root position does not further elucidate the biosynthetic substrate and product of the triterpenoid cyclase ancestor (i.e. whether it produced hopanoids, sterols, or a different molecule). Given that squalene-hopene cyclase was likely present in the common ancestor of Gracilicutes, one of the major divisions of bacteria (Santana-Molina et al., 2020), the OSC-SHC ancestor is likely quite ancient, although hopanes and steranes do not appear in the fossil record until 1.6 billion years ago (Brocks et al., 2005; French et al., 2015).

**Figure 7:**
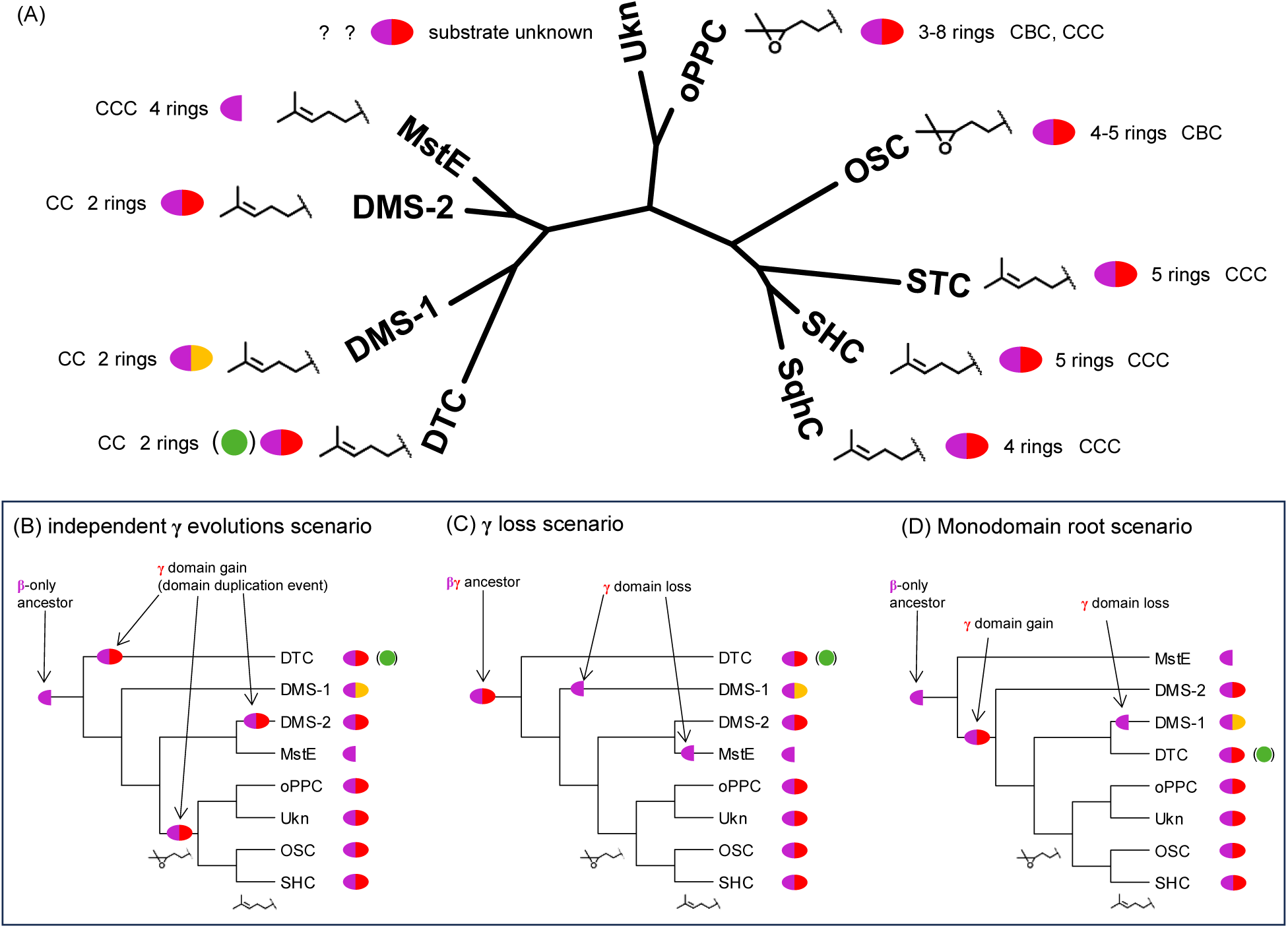
Distribution of terpenoid cyclase traits and hypotheses for superfamily root position. (A) Cyclase traits mapped across the unrooted tree, which has the same topology and internal branch lengths as the tree shown in Figure 2. Indicated traits include the number of rings in the final product, the structural conformation of the substrate (CBC: chair-boat-chair or CCC: chair-chair-chair); whether the substrate is epoxidized, and the number of domains (β = purple half-oval, γ = red half-oval, α (present in the plant subclade of diterpenoid cyclases) = green circle, haloacid dehalogenase domain = orange half-oval). (B-D) Alternate hypotheses for the root position of the cyclase tree with substrate and domain structure mapped. For simplicity, STC and SqhC are not shown, as their domain architecture and substrate functionalization are identical to SHC. OSC: oxidosqualene cyclases; SHC: squalene-hopene cyclases; MstE: meroterpenoid cyclases; DTC: diterpenoid cyclases; DMS: drimenyl synthases; Unk: homologs of unknown function; oPPC: epoxy-polyprenyl cyclases.

However, rooting on the diterpenoid cyclases implies either three independent evolutions of the γ domain (Figure 7B), or two losses of the γ domain on the MstE and DMS-1 stems (Figure 7C). The first scenario is not supported by the analysis of domain evolution discussed above (Figure 6). The second scenario is consistent with the domain trees but seems biochemically unlikely given that both domains are necessary for function in modern two-domain enzymes for substrate positioning and water exclusion (Abe, 2007; Fischer & Pearson, 2007; Wendt, 2005). The unusual cyclase DMS-1 may have had an easier path, with the novel haloacid dehalogenase-like domain (Vo et al., 2022) replacing the γ domain, or inserted between β and γ, in one mutational event, never requiring an evolutionary intermediate with a water-exposed active site. On the other hand, the monodomain cyclase merosterolic acid synthase (MstE) excludes water with a mobile loop that caps the active site when substrate is bound (Moosmann et al., 2020). This loop is significantly shorter in the closest related two-domain cyclase, drimenyl diphosphate synthase (Pan et al., 2022), suggesting that unless loop extension and domain loss occurred simultaneously, the active site of the MstE ancestor would have been exposed to water, making catalysis impossible. However, domain loss is not unknown in terpenoid cyclases (Oldfield & Lin, 2012). Biochemical investigation of basal or ancestral mono- and di-domain cyclases, as well as of, e.g., synthetic constructs of monodomain cyclases with shortened mobile loops, are necessary to establish the relative probabilities of domain loss and duplication in this superfamily.

If we then take domain loss in an MstE ancestor to be less likely than domain loss in a DMS-1 ancestor, a monodomain cyclase root (Figure 7D) becomes worth investigating. Rooting the structure-informed domain tree and the RMSD domain dendrogram on the monodomain cyclase (PDB ID 6SBB) places sesqui- and tri-terpenoid β domains as the next most basal, followed by remaining β domains, from which a monophyletic γ domain clade emerges. Although this position is in all cases far from the tree midpoint, it remains plausible given the evolutionary rarity of domain loss (Bridgham et al., 2009), and balances that rarity with the evidence for a single origin of the γ domain.

Within the triterpenoid cyclases, the oxidosqualene cyclase clade contains enzymes that form different numbers of rings (4 for sterol cyclases, 5 for arborinol cyclases). The squalene-hopene cyclases, which form 5 rings, have a stem group comprised of enzymes that form 4 (sporulenol cyclases) or 5 (squalene-tetrahymanol cyclase) rings respectively. Although we have found an outgroup with fewer rings (3 for Bra4 and PlaT2) as predicted by Fischer and Pearson (2007), this group also contains an enzyme that forms many more (8 for Lon15). A gradualist model of evolution, whereby additional rings evolve irreversibly and one-at-a-time, does not fit the data for this trait; rather, the number of rings is relatively labile. This is not surprising since anti-Markovnikov (secondary) carbocations are no longer considered to be cyclization intermediates (Hess, 2002; Hess & Smentek, 2013; Tantillo, 2010), removing the physical argument for a teleology of ring number.

Finally, substrate functionalization could have followed two evolutionary paths in the updated phylogeny. Fischer and Pearson (2007) hypothesized that the use of an epoxidized substrate evolved from an alkenyl ancestor once, on the oxidosqualene cyclase stem after the divergence of OSC and SHC. In the updated phylogeny, because epoxy-polyprenyl cyclases also used an epoxidized substrate, epoxide use could have evolved twice independently, once in the ancestor of each group. Because alkenyl-substrate enzymes can also cyclize epoxidized substrates, but epoxy-substrate enzymes cannot cyclize alkenyl substrates, evolutionary transitions from alkenyl to epoxy substrates seem in general more likely than the reverse. This is consistent with our discovery of enzymes within the squalene-hopene cyclase crown that only work on an epoxidized substrate (Table 1). Alternately, use of an epoxidized substrate may have evolved only once, in the common ancestor of oxidosqualene cyclases and epoxy-polyprenyl cyclases, and subsequently been lost in the squalene-hopene cyclase/squalene-tetrahymanol cyclase/sporulenol cyclase ancestor. Because epoxidized substrate biosynthesis requires molecular oxygen (Jahnke, 1986; Jahnke & Nichols, 1986), this scenario is less likely if the OSC-SHC-oPPC ancestor predates the radiation of Gracilicutes (Santana-Molina et al., 2020) and if Gracilicutes predate the rise of molecular oxygen around 2.4 billion years ago (Gumsley et al., 2017; Luo et al., 2016; Martinez-Gutierrez et al., 2023; Ostrander et al., 2024; Wang & Luo, 2025). However, it bears further investigation, e.g. via molecular clock and duplication-transfer-loss analysis, given the uncertainties in both antecedents. The relative probabilities of the two scenarios for substrate evolution will also be better-informed when the substrate of the “unknown” clade is discovered.

### Conclusions

Here we present an updated phylogenetic and functional analysis of triterpenoid cyclases. One result is that the triterpenoid cyclase active site has an elevated evolutionary rate on the SHC-STC-SqhC stem lineage. We hypothesize that this may be due to adaptation to an alkenyl substrate from an epoxy ancestor of all triterpenoid and epoxy-polyprenyl cyclases, *if* this ancestor evolved after the great oxidation event. Further, we found possible reversions to an epoxidized substrate in the SHC clade. This experimental result, as well as the discovery of a novel bacterial oxidosqualene-isoarborinol cyclase, shows that metagenomic cyclases from uncultured sources are a significant source of diversity and can help inform our understanding of the biochemical potential and evolutionary history of these fascinating enzymes. Future work focused on the functional and structural characterization of cyclases, particularly those from the uncharacterized clade sister to the epoxy-polyprenyl cyclases, will provide invaluable insight that will help constrain the evolutionary scenarios proposed in this study.

## Materials and Methods

### Gene synthesis and molecular cloning

Cyclase DNA sequences were codon-optimized for expression in *Escherichia coli* and synthesized through the Department of Energy Joint Genome Institute (DOE JGI) DNA Synthesis Science Program. These genes were obtained in the IPTG-inducible plasmid pSRKGm-*lac*UV5-rbs5 (Banta et al., 2017) in *E. coli* TOP10. Plasmid DNA was isolated using the GeneJET Plasmid Miniprep Kit (ThermoFisher Scientific) and sequenced to confirm promoter and gene sequences by ELIM Biopharm (Hayward, CA) using the following primers: 5′-AATGCAGCTGGCACGACAGG-3′ (forward) and 5′-CCAGGGTTTTCCCAGTCAC-3′ (reverse), purchased from Integrated DNA technologies (Coralville, IA).

### Bacterial culture and heterologous expression

*E. coli* DH10B expression strains were transformed by electroporation with the cyclase-bearing pSRKGm plasmid (Khan et al., 2008) and two other plasmids: pJBEI2997 (Addgene plasmid #351515) (Peralta-Yahya et al., 2011) encoding the MEV pathway and a pTrc99a derivative (Amann et al., 1988) encoding squalene synthase (and, in some cases, squalene monooxygenase). Gentamicin (15 μg/mL), carbenicillin (100 μg/mL), and/or chloramphenicol (20 μg/mL) were added to media as appropriate to select for maintenance of each plasmid. Plasmids are described in detail in Supplementary Table 4.

Strains were cultured in 50 mL TYGPN broth (RECIPE) at 37 °C while shaking at 225 rpm. Cultures were induced with 500 μM isopropyl *β*-D-1-thiogalactopyranoside (IPTG) at an OD_600_ of ∼0.6, then incubated an additional 48 h at 30 °C while shaking at 225 rpm. Cells were harvested by centrifugation at 4500 × g for 10 min at 4 °C. Cell pellets were stored at −20 °C until lipid extraction. In every experiment, a negative control strain with an empty pSRKgm plasmid and relevant positive controls (strains with pSRKGm containing a triterpenoid cyclase of known function) were grown and extracted alongside experimental strains.

### Lipid extraction

Cell pellets were extracted using a modified Bligh–Dyer method (Bligh & Dyer, 1959; Welander et al., 2012): cells were resuspended in 2 mL of deionized water and transferred to a solvent-washed Teflon centrifuge tube containing 5 mL of methanol and 2.5 mL of dichloromethane (DCM), vortexed for 30 seconds, and sonicated for 1 hour in a water bath sonicator. 10 mL of deionized water and 10 mL of DCM were then added, and samples were vortexed and incubated overnight at −20 °C. Samples were centrifuged for 10 min at 2,800 × g, and the organic layer was transferred to a baked glass vial and evaporated at 40 °C under a gentle stream of N_2_ to yield the total lipid extract (TLE). TLE was stored at −20 °C, and derivatized to trimethylsilyl ethers with 1:1 (v:v) bis(trimethylsilyl)trifluoroacetamide (BSTFA):pyridine for 1 h at 70 °C before analysis by gas chromatography–mass spectrometry (GC-MS). The alcohol-soluble fractions of some TLEs were further purified by silicon column chromatography before derivatization, using a solvent series of hexanes, 8:2 hexanes:DCM, DCM, 1:1 DCM:ethyl acetate, and ethyl acetate (Summons et al., 2013).

### GC-MS analysis

Lipids were separated with an Agilent 7890B Series GC equipped with two Agilent DB-17HT columns (30 m × 0.25 mm i.d. × 0.15 μm film thickness) in tandem with helium as the carrier gas at a constant flow of 1.1 ml/min and programmed as follows: 100 °C for 2 min, then 12 °C/min to 250 °C and held for 10 min, then 10 °C/min to 330 °C and held for 17.5 min. 2 μL of each sample was injected in splitless mode at 250 °C. The GC was coupled to an Agilent 5977 A Series MSD with the ion source at 230 °C and operated at 70 eV in electron ionization (EI) mode scanning from 50 to 850 Da in 0.5 s. All lipids except isoarborinol were identified based on their retention time and comparison with published spectra. Isoarborinol was identified as a likely sterol by GC-MS, and its structure was confirmed by nuclear magnetic resonance spectroscopy (NMR).

### NMR Analysis

NMR experiments were performed following Banta et al. (2017). TLE saponification was accomplished by heating with 10% (vol/vol) sodium hydroxide (NaOH)/MeOH at reflux for 16 h. The reaction mixture was partitioned between water and hexane/ethyl acetate (EtOAc) 2:1; the organic layers were filtered through neutral alumina and concentrated to dryness with a stream of nitrogen. The saponified lipids were fractionated by preparative TLC on glass-backed plates (10 cm in length) coated with a 0.25-mm layer of silica gel 60 F254 using hexane/EtOAc 4:1 as the developing solvent. The triterpenol fraction was further fractionated by reversed-phase HPLC with a system consisting of a Waters 6000A pump, Waters 410 differential refractometer, and two Altex Ultrasphere ODS 5-μm 10 × 250 mm columns in series using a flow rate of 3 mL/min MeOH. After evaporation of the HPLC solvent, the triterpenols were characterized by NMR using a Bruker Avance III HD with an Ascend 800 MHz magnet and a 5-mm TCI cryoprobe at 30 °C using deuterated chloroform (CDCl3) as the solvent. Calibration was by the residual solvent signal (^1^H: 7.26 ppm; ^13^C 77.0 ppm).

### Sequence search and curation

Cyclase homologs were retrieved from the Joint Genome Institute Integrated Microbial Genomes & Microbiomes database (JGI IMG; https://img.jgi.doe.gov) and the National Center for Biological Informatics nonredundant sequence database (https://www.ncbi.nlm.nih.gov/genbank) using the set of query sequences listed in Supplementary Table 5. Protein-protein BLAST algorithms were used in both cases, with an expect threshold of 0.05 and a word size of 5 for NCBI, and an e-value cutoff of 1e-5 for JGI. Additionally, we downloaded all hits for Pfams PF13243 (squalene-hopene cyclase C-terminal domain) and PF13249 (squalene-hopene cyclase N-terminal domain) from the European Bioinformatics Institute (EBI; https://www.ebi.ac.uk/interpro/entry/pfam/#table). Combined, these searches resulted in an initial set of 44,209 unique sequences. BLAST results were filtered by length dependent on query sequence as follows: 600-800 amino acids (aa) for triterpenoid cyclases, 550-700 aa for epoxy-polyprenyl cyclases and members of the “unknown function” clade, 700-1,000 aa for plant and fungal diterpenoid cyclases, 450-600 aa for bacterial diterpenoid cyclases, 550-650 aa for DMS1, 450-550 aa for DMS2, and 300-400 aa for MstE. These length ranges were determined based on length distribution of results and length range of functionally characterized representatives of these groups. Sequences from EBI were filtered to a length range of 350 (the length of the smallest known Type II cyclase; Moosmann et al., 2020) to 900 (the length of the largest known Type II domain-bearing cyclase; Mitsuhashi et al., 2017) amino acids. Remaining sequences were subsetted with CD-hit (command cd-hit -i in.fasta -o out.fasta -c 0.90 -n 5 -M 6000 -d 0 -T 8) (Fu et al., 2012) and aligned with the MAFFT v7.402 fftns2 algorithm (Rozewicki et al., 2019), after which sequences lacking an aligned DxD motif were removed (except for oPPC homologs with an [E/D]SA[E/N] motif; Dürr et al., 2006; Hayashi et al., 2008) to yield a database of 13,680 sequences.

### Phylogenetic estimation

All sequences in the database described above were aligned using the MAFFT linsi algorithm (Rozewicki et al., 2019) implemented in Magus (Smirnov & Warnow, 2021). The alignment was trimmed using trimAl with a gap threshold of 0.1 (command trimal -in in.aln -out out.aln -gt 0.1), and subsetted with a maximum sequence identity of 50% (command trimal -in in.aln -out out.aln -maxidentity 0.5) (Capella-Gutiérrez et al., 2009). Model selection and maximum likelihood phylogeny estimation were performed with IQ-TREE 2.2.0 (Bui et al., 2020), with the model EX_EHO+R5 (Le & Gascuel, 2010) found to be the best model by Bayesian Information Criterion and Akaike Information Criterion (Kalyaanamoorthy et al., 2017). For all phylogeny estimation, we performed ≤ 3,000 ultrafast bootstrap replicates (Hoang et al., 2018). All input commands are reported in Supplementary Table 2. Resulting phylogenetic trees were visualized in Dendroscope (Huson & Scornavacca, 2012), Figtree (Rambaut, 2009), or iTOL (Letunic & Bork, 2021).

### Structural phylogenetics

We used the rigid-body superposition algorithm mTM-align (R. Dong et al., 2018) to align representative cyclase crystal structures (Supplementary Table 6). To generate the character state matrix, we used this multiple structure alignment to measure and encode every chemical and structural character of the proteins that was both unlikely to be subject to artifacts of crystallization and likely to evolve as slowly or slower than the protein’s amino acid sequence: reaction class, active site motif, subunits, loops and helices, and N-terminal tail. The resulting character state matrix became input for Bayesian phylogeny estimation using MrBayes 3.2.6 (Ronquist et al., 2012). We compared these phylogenies to LDDT and TM score trees calculated using a neighbor-joining algorithm implemented in Foldtree (Moi et al., 2023), as well as structural trees calculated using the FoldTree method and sequence trees.

### Analysis of adaptation

Using the phylogeny estimated under the model EX_EHO+R5 in IQ-TREE, ancestral protein sequences were reconstructed using IQ-TREE. For each of the branches between major clades (highlighted in Figure 5), most-likely ancestral sequences at the branches’ terminal nodes were modeled using ColabFold (Mirdita et al., 2022). We then used Fisher’s exact test (2-tail, multiple-test corrected) to measure whether the amino acid substitutions that occurred along that branch (differences between the two ancestral proteins) are statistically clustered in functional regions of the protein. 6 functional regions were tested: the protein’s N terminus, the C terminus, the helix that acts as a membrane anchor, the dimer contact surface, the tunnel from the surface to the active site, and the active site itself (precise region definitions can be found in Supplementary Table 7). Substrate tunnel residues were calculated with Caver (Pavelka et al., 2016) and active site was defined as residues within 5 Å of substrate molecule.

## Data availability

The data supporting the findings in this study are available in the main text and supplementary information. The terpenoid cyclase alignment and phylogeny (shown in Figures 2, 3, and 5) are available as supplementary files.

## Acknowledgments

The authors would like to thank the students of ESS 143: Molecular Geomicrobiology Lab for hard work and curiosity in initial experiments characterizing the lipid products of several metagenomic triterpenoid cyclases. The authors would like to thank members of the Welander lab, M. Horst, B. Kapili, C.D. Stark, and B. Barros-McShea for critically examining, and thereby improving, this work. We would also like to thank Stanford Research Computing for access to and assistance with the Sherlock High-Performance Computing Environment, where we performed phylogeny estimation and protein structure prediction, and the Stanford Geomicrobiology Shared Laboratories Core Facility (RRID:SCR_025000), where we performed portions of the experimental work. H.S.M. and P.V.W. were supported by NSF Grant EAR-1752564. H.S.M. was supported by the National Science Foundation (NSF) Graduate Research Fellowship, the Stanford Enhancing Diversity in Graduate Education (EDGE) Fellowship, and the Stanford School of Sustainability McGee and Levorsen Graduate Research Grant. R.A.V. was supported by the Stanford Earth Summer Undergraduate Research (SESUR) Program. B.O. and J.L.G. were supported (in part) by National Institutes of Health (NIH) Grant R15GM143714; the 800 MHz NMR was acquired through NIH S10 OD012254.

