## Supplemental Materials for "The evolutionary history and modern diversity of triterpenoid cyclases"

### Supplementary Information for McShea et al.,
#### "*The evolutionary history and modern diversity of triterpenoid cyclases*"

#### Supplementary Figures

##### **Supplementary Figure 1:** Representative gas chromatograms.

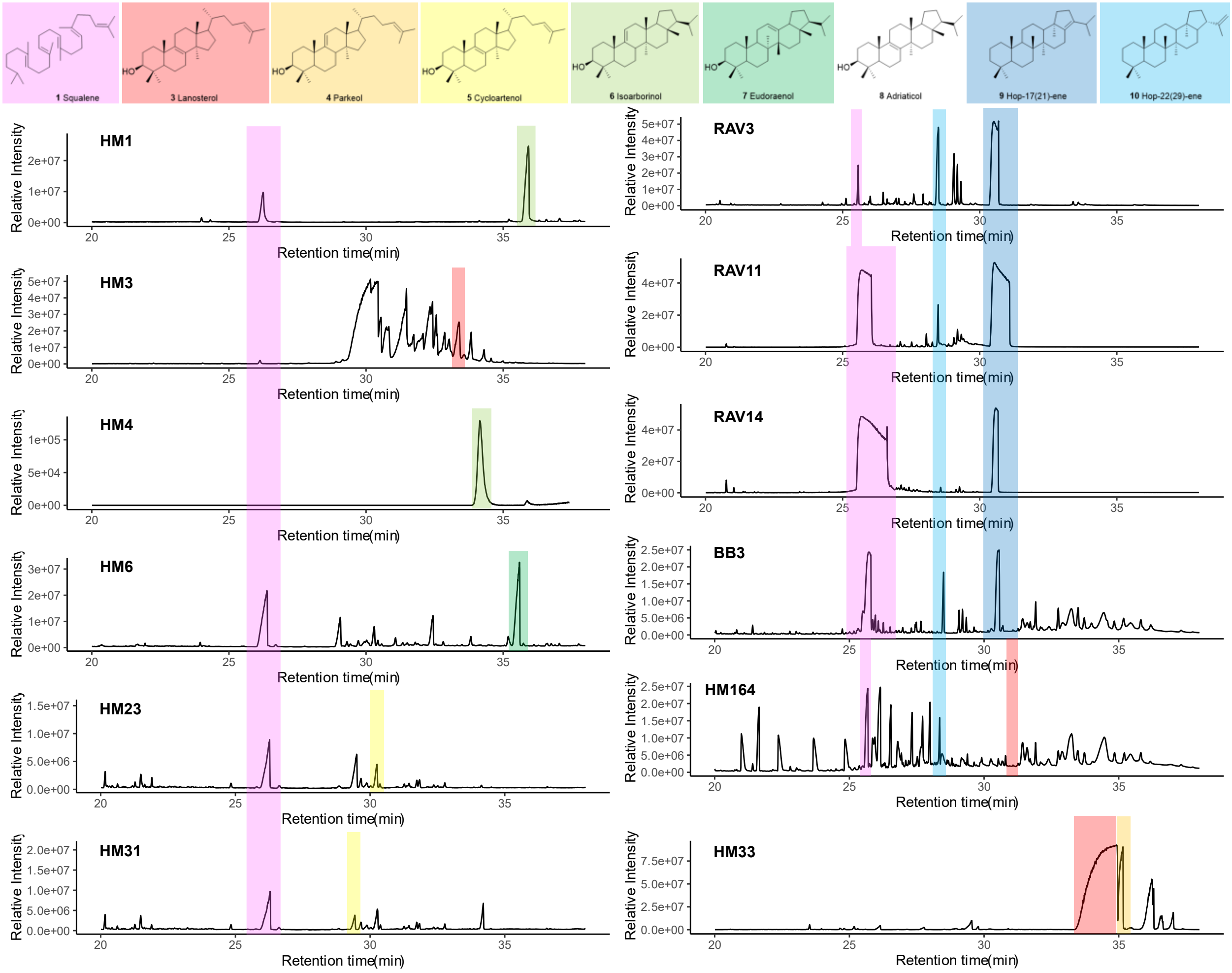

**Supplementary Figure 2:** Representative mass spectra.

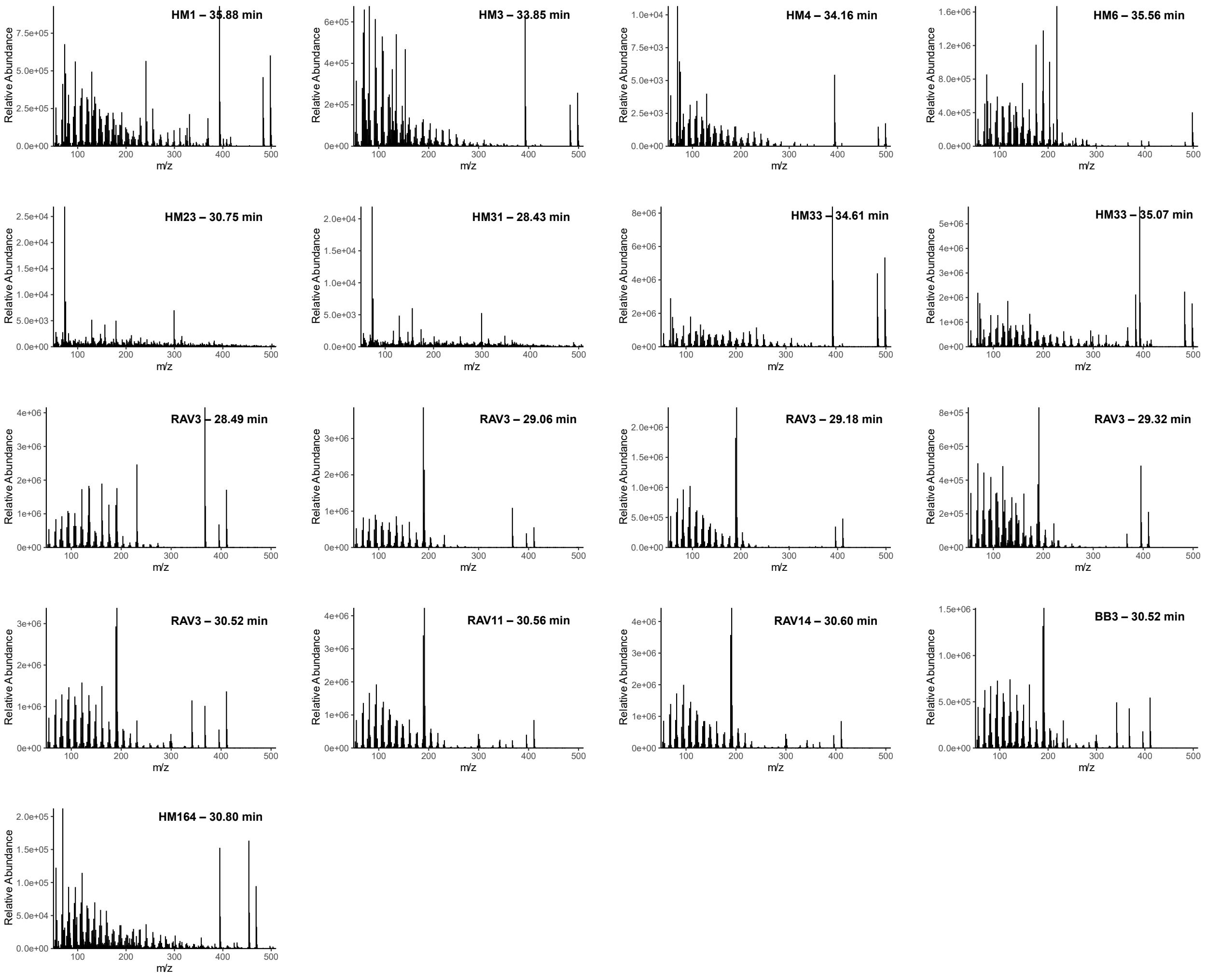

**Supplementary Figure 3:** Raw NMR spectra of isoarborinol. Panels in order of appearance: (A) 800 MHz 1H-NMR spectrum. (B) 201 MHz 13C-NMR spectrum. (C) 2D NMR (HSQC-DEPT) spectrum. (D) 2D NMR (HMBC) spectrum. Chemical shifts are summarized in Supplementary Table 3.

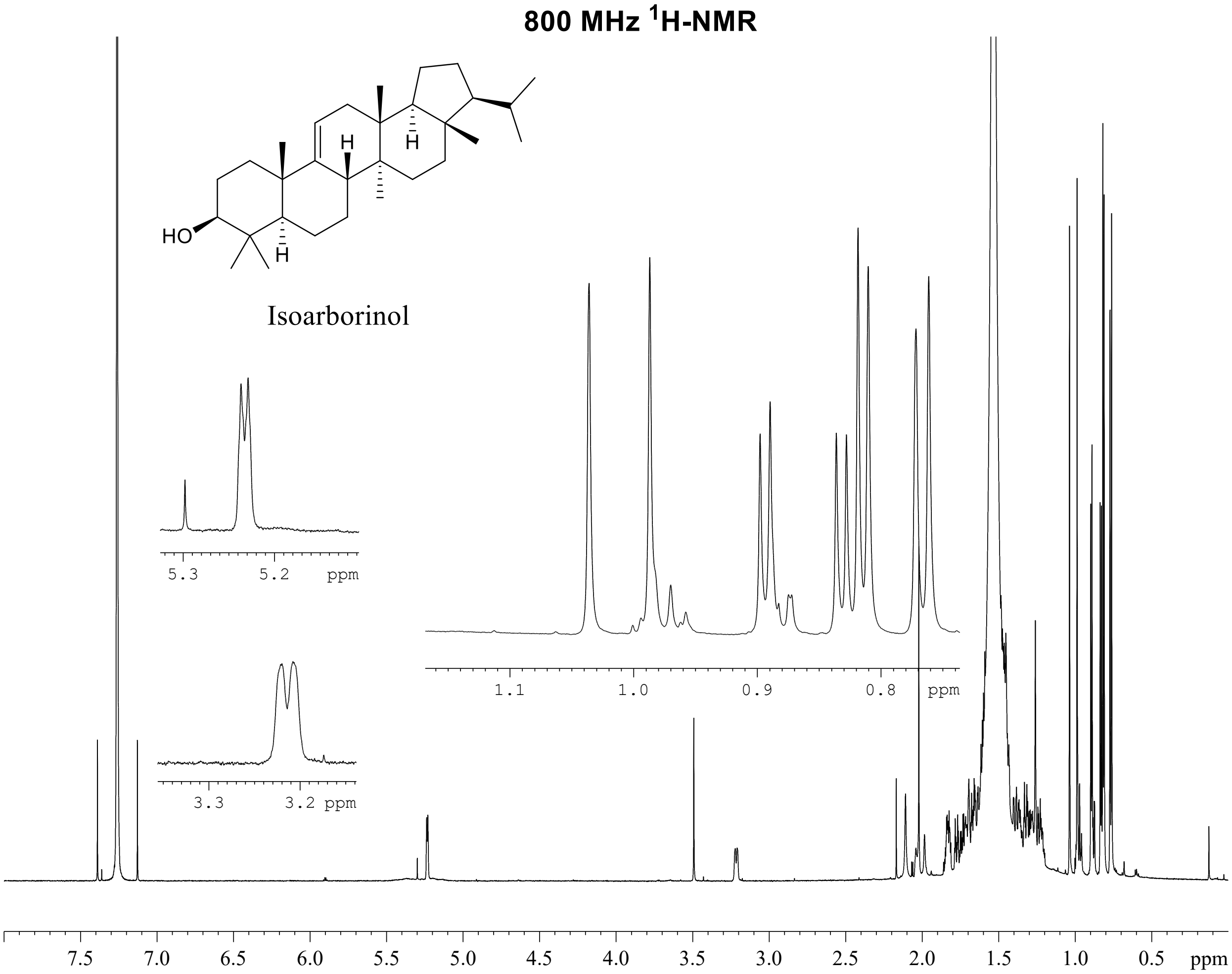

**201 MHz $^{13}$C-NMR**

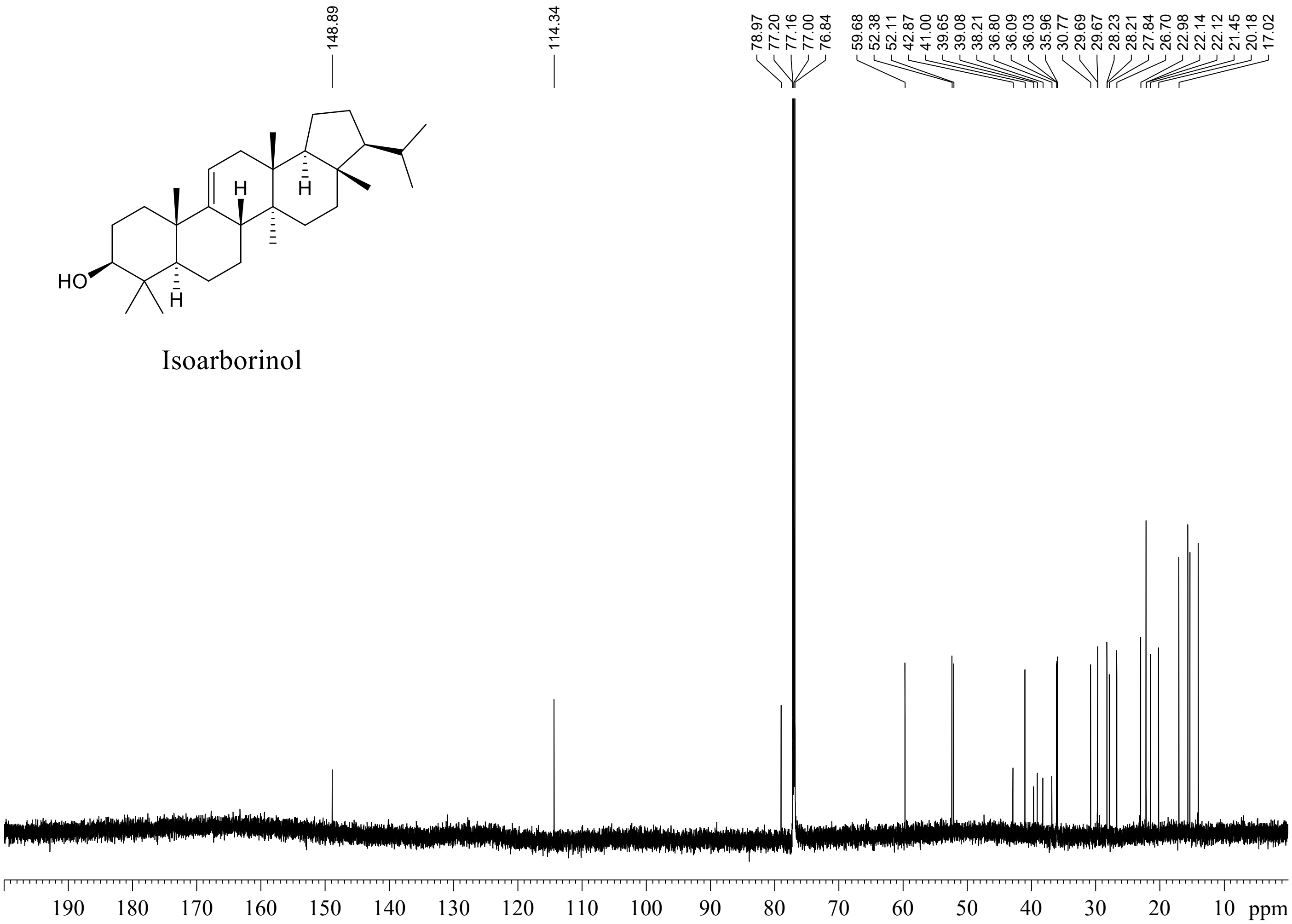

**800 MHz HSQC-DEPT**

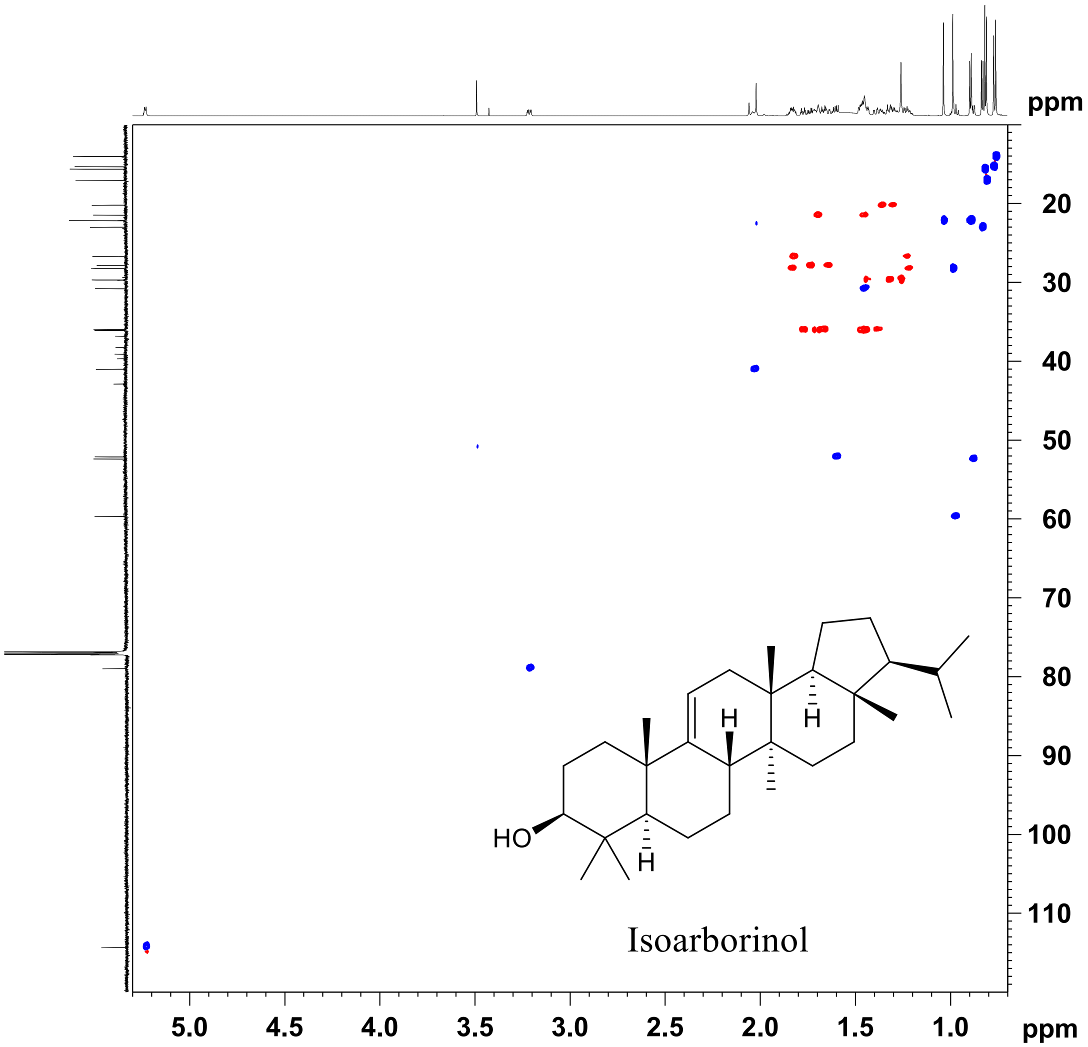

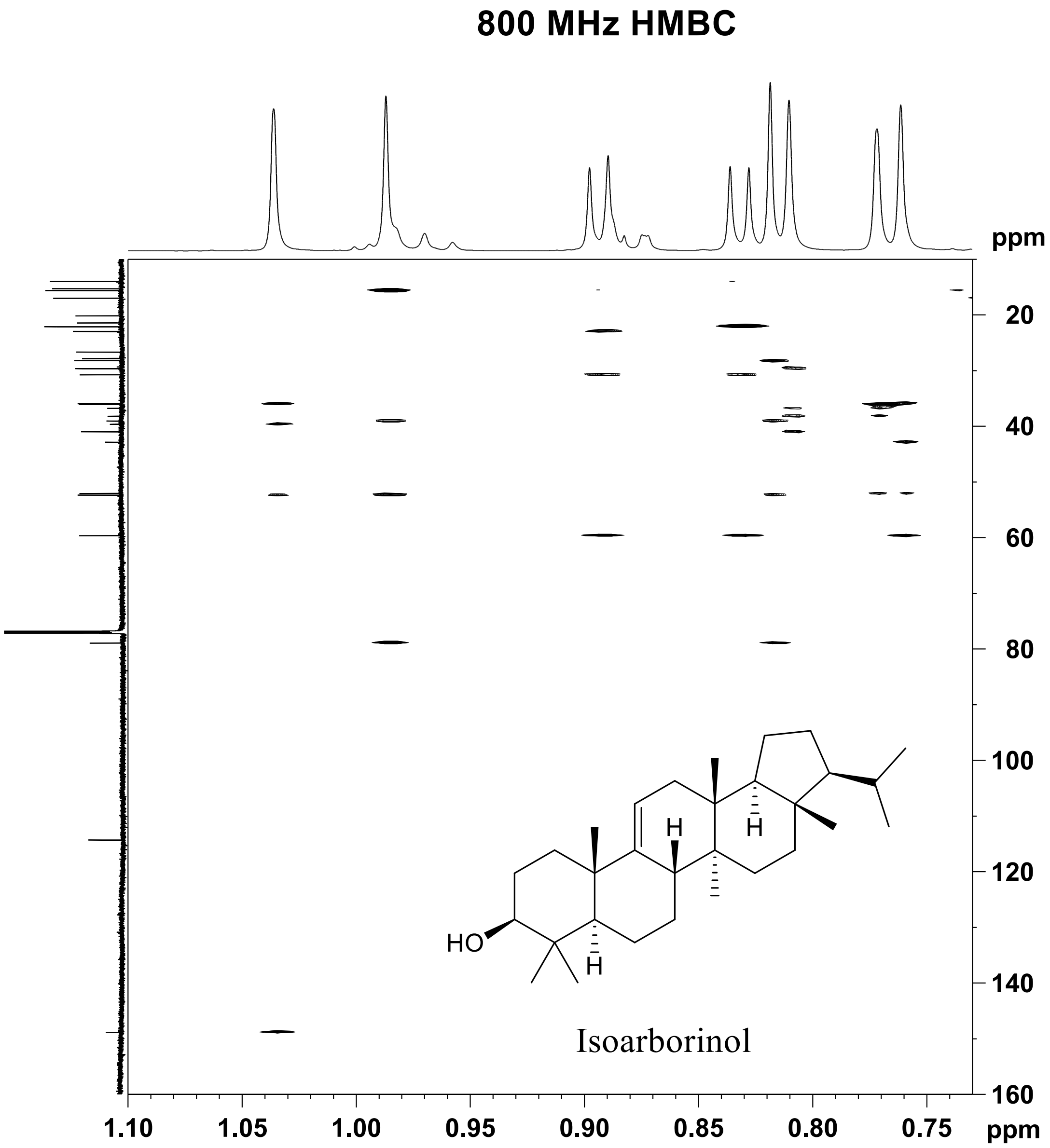


**Supplementary Figure 3:**

**Supplementary Tables (all tables are also in Supplement.xlsx)**

**Supplementary Table 1:** Pairwise root mean squared deviation (RMSD) of terpenoid cyclase β and γ domain crystal structures, labeled by PDB ID.

| | 1dce-β | 1fpp-β | 1n1b-β | 1n4p-β | 1w6k-β | 1w6k-γ | 2j5c-β | 2ong-β | 2sqc-β | 2sqc-γ | 3g4d-β | 3n0f-β | 3p5p-β | 3p5p-γ | 3pya-β | 3pya-γ | 3s9v-β | 3s9v-γ | 3sae-β | 3sae-γ | 4fjq-β | 5bp8-β | 5bp8-γ | 5c05-β | 5eas-β | 5hlr-β | 5uv0-β | 6j74-β | 6sbb-β | 6vpt-β | 6vpt-γ | 7xkx-β | 7xkx-γ | 7xq4-β |
|---|---|---|---|---|---|---|---|---|---|---|---|---|---|---|---|---|---|---|---|---|---|---|---|---|---|---|---|---|---|---|---|---|---|---|
| 1fpp-β | 2.1 | | | | | | | | | | | | | | | | | | | | | | | | | | | | | | | | | |
| 1n1b-β | 3.9 | 4.2 | | | | | | | | | | | | | | | | | | | | | | | | | | | | | | | | |
| 1n4p-β | 1.7 | 2.0 | 4.4 | | | | | | | | | | | | | | | | | | | | | | | | | | | | | | | |
| 1w6k-β | 3.5 | 4.8 | 4.5 | 4.1 | | | | | | | | | | | | | | | | | | | | | | | | | | | | | | |
| 1w6k-γ | 4.4 | 4.9 | 4.7 | 4.7 | 4.6 | | | | | | | | | | | | | | | | | | | | | | | | | | | | | |
| 2j5c-β | 4.1 | 4.1 | 1.4 | 4.2 | 4.0 | 5.4 | | | | | | | | | | | | | | | | | | | | | | | | | | | | |
| 2ong-β | 4.1 | 4.2 | 1.2 | 4.4 | 4.8 | 5.6 | 1.5 | | | | | | | | | | | | | | | | | | | | | | | | | | | |
| 2sqc-β | 3.7 | 3.9 | 4.7 | 3.9 | 2.7 | 4.9 | 4.6 | 4.5 | | | | | | | | | | | | | | | | | | | | | | | | | | |
| 2sqc-γ | 4.5 | 4.9 | 5.6 | 4.3 | 4.6 | 2.4 | 5.2 | 5.6 | 4.9 | | | | | | | | | | | | | | | | | | | | | | | | | |
| 3g4d-β | 3.8 | 4.1 | 1.6 | 4.2 | 4.3 | 5.7 | 1.4 | 1.9 | 4.4 | 5.7 | | | | | | | | | | | | | | | | | | | | | | | | |
| 3n0f-β | 3.8 | 4.5 | 2.2 | 4.4 | 4.4 | 5.5 | 1.3 | 2.2 | 4.5 | 4.6 | 1.4 | | | | | | | | | | | | | | | | | | | | | | | |
| 3p5p-β | 4.2 | 4.2 | 2.6 | 4.1 | 3.9 | 4.4 | 2.5 | 2.6 | 4.2 | 4.4 | 2.2 | 2.4 | | | | | | | | | | | | | | | | | | | | | | |
| 3p5p-γ | 4.3 | 4.3 | 4.7 | 4.6 | 4.2 | 4.3 | 4.5 | 5.0 | 3.9 | 3.9 | 4.6 | 4.9 | 4.5 | | | | | | | | | | | | | | | | | | | | | |
| 3pya-β | 4.5 | 4.5 | 2.3 | 4.5 | 3.9 | 4.6 | 2.2 | 2.6 | 4.3 | 4.7 | 2.0 | 2.4 | 1.4 | 4.7 | | | | | | | | | | | | | | | | | | | | |
| 3pya-γ | 4.4 | 4.3 | 4.6 | 4.4 | 4.1 | 4.0 | 4.8 | 4.7 | 3.7 | 3.9 | 4.8 | 5.2 | 4.4 | 2.0 | 4.5 | | | | | | | | | | | | | | | | | | | |
| 3s9v-β | 4.5 | 4.5 | 2.6 | 4.5 | 4.1 | 4.2 | 2.2 | 2.6 | 4.3 | 4.4 | 2.0 | 2.4 | 1.2 | 4.2 | 1.5 | 4.3 | | | | | | | | | | | | | | | | | | |
| 3s9v-γ | 4.4 | 4.7 | 4.8 | 4.7 | 4.0 | 4.0 | 4.8 | 4.4 | 4.1 | 4.1 | 4.7 | 4.6 | 4.5 | 1.8 | 4.7 | 1.3 | 4.3 | | | | | | | | | | | | | | | | | |
| 3sae-β | 4.3 | 4.6 | 2.0 | 4.1 | 4.4 | 4.6 | 2.3 | 2.2 | 4.9 | 4.5 | 2.6 | 2.4 | 2.1 | 4.8 | 2.4 | 4.5 | 2.4 | 4.7 | | | | | | | | | | | | | | | | |
| 3sae-γ | 4.3 | 4.4 | 4.8 | 4.4 | 4.2 | 4.3 | 4.7 | 5.0 | 4.1 | 4.4 | 4.6 | 4.8 | 4.4 | 2.0 | 4.5 | 1.6 | 4.5 | 1.4 | 4.5 | | | | | | | | | | | | | | | |
| 4fjq-β | 5.0 | 4.5 | 2.4 | 4.5 | 4.4 | 5.4 | 2.2 | 2.4 | 4.9 | 5.8 | 2.0 | 2.4 | 2.3 | 4.8 | 2.6 | 4.7 | 2.5 | 4.7 | 2.9 | 4.9 | | | | | | | | | | | | | | |
| 5bp8-β | 3.5 | 3.6 | 4.3 | 3.7 | 2.7 | 4.0 | 4.1 | 4.3 | 2.9 | 5.2 | 4.0 | 4.1 | 3.6 | 4.0 | 3.7 | 4.3 | 3.6 | 4.5 | 4.1 | 4.6 | 4.6 | | | | | | | | | | | | | |
| 5bp8-γ | 4.5 | 4.9 | 4.5 | 4.6 | 4.4 | 4.8 | 4.6 | 5.1 | 5.2 | 4.8 | 4.7 | 4.9 | 5.2 | 2.7 | 4.8 | 3.1 | 4.1 | 2.9 | 4.8 | 3.0 | 5.0 | 4.9 | | | | | | | | | | | | |
| 5c05-β | 3.8 | 3.9 | 1.1 | 4.6 | 4.6 | 4.7 | 1.6 | 1.3 | 5.0 | 5.5 | 1.8 | 2.2 | 2.2 | 5.0 | 2.0 | 4.8 | 2.1 | 4.9 | 2.0 | 4.8 | 2.2 | 4.3 | 5.0 | | | | | | | | | | | |
| 5eas-β | 4.1 | 4.0 | 1.8 | 4.2 | 4.4 | 5.3 | 1.4 | 1.9 | 4.7 | 5.4 | 1.4 | 1.7 | 2.2 | 4.3 | 2.1 | 4.9 | 2.2 | 4.9 | 2.5 | 4.8 | 1.9 | 4.1 | 4.7 | 1.8 | | | | | | | | | | |
| 5hlr-β | 4.2 | 4.3 | 4.8 | 4.5 | 4.1 | 5.5 | 4.5 | 4.6 | 4.1 | 5.3 | 4.5 | 4.6 | 4.4 | 4.4 | 4.4 | 4.7 | 4.6 | 4.1 | 5.0 | 4.6 | 5.2 | 3.5 | 4.6 | 4.9 | 4.7 | | | | | | | | | |
| 5uv0-β | 4.0 | 4.4 | 2.0 | 4.4 | 4.4 | 5.5 | 1.9 | 1.9 | 4.8 | 5.8 | 2.0 | 2.2 | 2.3 | 5.0 | 2.2 | 5.0 | 2.3 | 5.1 | 2.3 | 4.9 | 2.5 | 4.2 | 4.9 | 2.0 | 2.1 | 4.4 | | | | | | | | |
| 6j74-β | 0.8 | 2.1 | 3.8 | 1.7 | 3.4 | 4.4 | 4.1 | 4.0 | 3.8 | 4.5 | 4.2 | 3.8 | 4.3 | 4.3 | 4.6 | 4.5 | 4.5 | 4.4 | 4.2 | 4.4 | 4.4 | 3.7 | 4.5 | 3.8 | 4.3 | 4.4 | 4.1 | | | | | | | |
| 6sbb-β | 3.9 | 3.8 | 4.9 | 3.9 | 3.0 | 4.1 | 4.2 | 4.5 | 3.1 | 3.9 | 4.4 | 4.5 | 3.9 | 3.9 | 4.0 | 3.9 | 3.6 | 4.1 | 4.7 | 4.2 | 4.5 | 2.4 | 4.3 | 4.6 | 4.0 | 4.0 | 4.3 | 3.8 | | | | | | |
| 6vpt-β | 3.8 | 3.7 | 4.1 | 3.8 | 2.9 | 4.8 | 4.3 | 4.6 | 2.7 | 4.6 | 4.4 | 4.6 | 3.8 | 4.3 | 3.8 | 4.5 | 4.0 | 4.3 | 4.5 | 4.8 | 4.5 | 2.3 | 4.8 | 4.2 | 4.4 | 3.8 | 4.6 | 3.8 | 2.7 | | | | | |
| 6vpt-γ | 4.3 | 4.3 | 4.7 | 4.7 | 3.8 | 3.8 | 4.4 | 4.6 | 3.7 | 4.1 | 4.7 | 4.9 | 4.1 | 2.3 | 4.4 | 2.0 | 4.3 | 2.1 | 4.5 | 2.1 | 4.8 | 4.9 | 2.7 | 4.9 | 4.5 | 4.7 | 4.8 | 4.3 | 4.2 | 4.6 | | | | |
| 7xkx-β | 3.5 | 3.5 | 4.3 | 3.6 | 2.7 | 5.0 | 4.0 | 4.2 | 3.0 | 5.0 | 4.0 | 3.9 | 3.6 | 4.4 | 3.6 | 4.4 | 3.5 | 4.2 | 4.2 | 4.5 | 4.5 | 1.9 | 5.0 | 4.2 | 3.9 | 3.5 | 4.1 | 3.6 | 2.5 | 2.2 | 4.6 | | | |
| 7xkx-γ | 4.4 | 4.4 | 4.8 | 5.0 | 4.2 | 4.2 | 4.9 | 5.0 | 4.2 | 4.1 | 4.6 | 5.0 | 4.4 | 2.8 | 4.8 | 3.1 | 4.1 | 3.1 | 4.4 | 3.2 | 5.1 | 4.6 | 2.2 | 4.8 | 4.9 | 4.6 | 4.4 | 4.5 | 4.0 | 4.9 | 3.1 | 4.3 | | |
| 7xq4-β | 3.6 | 3.7 | 4.6 | 3.9 | 2.2 | 4.2 | 3.7 | 4.8 | 2.0 | 4.3 | 4.0 | 3.9 | 4.0 | 3.7 | 3.9 | 3.7 | 4.0 | 3.8 | 4.7 | 3.9 | 4.8 | 2.5 | 4.2 | 3.9 | 4.0 | 3.5 | 4.6 | 3.7 | 2.4 | 2.6 | 4.6 | 2.8 | 3.9 | |
| 7xq4-γ | 4.3 | 4.3 | 4.7 | 4.8 | 3.8 | 3.2 | 4.6 | 4.8 | 4.5 | 3.4 | 4.9 | 4.8 | 4.4 | 3.7 | 4.4 | 4.1 | 4.3 | 4.2 | 5.1 | 4.2 | 4.6 | 4.6 | 4.3 | 4.8 | 4.7 | 4.7 | 4.8 | 4.8 | 3.5 | 4.4 | 3.6 | 4.3 | 4.4 | 4.5 |

**Supplementary Table 2:** IQ-TREE input parameters for all phylogeny estimations in this study. The tree estimated under EX_EHO+R5 and its input alignment are available as supplementary newick and fasta files.

| **Tree** | **Command** | **Gross topology** |
|---|---|---|
| EX_EHO | iqtree2 -s in-50.fasta -m EX_EHO+R5 -B 1000 -bnni -T AUTO | As shown in Figure 2 |
| Q.pfam | iqtree2 -s in-50.fasta -m Q.pfam+R5 -B 1000 -bnni -T AUTO | As shown in Figure 2 |
| β domain only | iqtree2 -s in-50-Bonly.fasta -m EX_EHO+R5 -B 1000 -bnni -T AUTO | As shown in Figure 2 |
| γ domains aligned separately | iqtree2 -s in-50-gg.fasta -m EX_EHO+R5 -B 1000 -bnni -T AUTO | As shown in Figure 2 |
| Untrimmed alignment | iqtree2 -s in-50-untr.fasta -m EX_EHO+R5 -B 1000 -bnni -T AUTO | As shown in Figure 2 |
| 70% identity cutoff | iqtree2 -s in-70.fasta -m EX_EHO+R5 -B 1000 -bnni -T AUTO | As shown in Figure 2, with more sequences |
| Figure 6 sequence tree | iqtree2 -s struct.fasta -m Q.pfam+F+G4 -B 1000 -nt AUTO | As shown in Figure 6 |

**Supplementary Table 3:** 800 Mz $^{1}H$ and 201 Mz $^{13}C$ NMR assignments. All $^{13}C$ data are in boldface; coupling constants are given in Hz. The assignments were determined by HSQC-DEPT, HMBC and COSY 2D NMR experiments. $^{1}H$ and $^{13}C$ chemical shifts are given to three and two decimal places, respectively, if directly observed; two and one decimal places, if determined from 2D spectra.

| Position | Isoarborinol |
|---|---|
| 1 | **36.03**<br>1.774 dt 13.0, 3.5; 1.46 |
| 2 | **27.84**<br>1.74; 1.64 |
| 3 | **78.97**<br>3.214 dd 11.8, 4.3 |
| 4 | **39.08** |
| 5 | **52.38**<br>0.88 |
| 6 | **21.45**<br>1.70; 1.46 |
| 7 | **26.70**<br>1.83; 1.23 |
| 8 | **41.00**<br>2.03 |
| 9 | **148.89** |
| 10 | **39.65** |
| 11 | **114.34**<br>5.233 dt 6.1, 1.6 |
| 12 | **36.09**<br>1.71; 1.46 |
| 13 | **36.80** |
| 14 | **38.21** |
| 15 | **29.69**<br>1.45; 1.32 |
| 16 | **35.96**<br>1.667 dt 12.9, 3.2; 1.385 dt 3.1, 13.1 |
| 17 | **42.87** |
| 18 | **52.11**<br>1.601 dd 13.0, 7.1 |
| 19 | **20.18**<br>1.36; 1.31 |
| 20 | **28.21**<br>1.84; 1.22 |
| 21 | **59.68**<br>0.977 q 9.8 |
| 22 | **30.77**<br>1.46 |
| 23 | **28.23**<br>0.987, s |
| 24 | **15.62**<br>0.819, s |
| 25 | **22.14**<br>1.036, s |
| 26 | **17.02**<br>0.810, s |
| 27 | **15.30**<br>0.772, s |
| 28 | **14.00** |

| | |
|---|---|
| | 0.761, s |
| 29 | **22.98**<br>0.832, d 6.5 |
| 30 | **22.12**<br>0.894, d 6.6 |

**Supplementary Table 4:** All plasmids used in this study.

| **Plasmid** | **Description** | **Source** |
|---|---|---|
| pJBEI2997 (pABB302) | p15A ori, $Cm^R$, Addgene #35151 (pBbA5c-MevT(CO)-MBIS(CO, ispA)), J. Keasling and T.S. Lee. Expresses the MEV pathway to produce farnesyl pyrophosphate. | (Peralta-Yahya et al., 2011) |
| pTrc-sqs (pABB303) | pTrc99a derivative with pBR322 ori, *lac*UV5 promoter, $Amp^R$. Expresses *Methylomicrobium alcaliphilum* MEALZ_3096 (squalene synthase). | (Amann et al., 1988; Banta et al., 2017) |
| pTrc-sqs-synRBS-smo (pABB497) | pTrc99a derivative with pBR322 ori, *lac*UV5 promoter, $Amp^R$, stronger ribosome binding site. Expresses *Methylomicrobium alcaliphilum* MEALZ_3096-MEALZ_0767 (squalene synthase-squalene epoxidase). | (Amann et al., 1988; Banta et al., 2017) |
| pSRKGm-lacUV5-rbs5 (pABB492) | pSRK derivative with pBBR1 ori, *lac*UV5 promoter, $Gm^R$, stronger ribosome binding site from pABB251. All cyclases were cloned into this plasmid. | (Banta et al., 2017; Khan et al., 2008) |

**Supplementary Table 5:** Query sequences for database construction, representing the known functional and phylogenetic diversity of terpenoid cyclases.

| Taxon | Species | Substrate | Architecture | Length | Short name | Accession |
|---|---|---|---|---|---|---|
| Bacteria | Bradyrhizobium diazoefficiens USDA 110 | C20 | BG | 587 | BjCPS | BAC47414.1 |
| Bacteria | Nocardia terpenica | C20 | BG | 556 | Bra4 | BAG16278.1 |
| Bacteria | Kitasatospora griseola | C20 | BG | 499 | Cyc1 | BAB39206.1 |
| Bacteria | Streptomyces sp. KO-3988 | C20 | BG | 511 | Orf2 | BAD86797.1 |
| Bacteria | Streptomyces sp. Tu6071 | C20 | BG | 571 | PlaT2 | ABB69743.1 |
| Bacteria | Streptomyces platensis | C20 | BG | 533 | PtmT2 | ACO31276.1 |
| Bacteria | Streptomyces platensis | C20 | BG | 533 | PtnT2 | ADD83015.1 |
| Bacteria | Mycobacterium tuberculosis H37Rv | C20 | BG | 501 | Rv3377c | NP_217894.1 |
| Bacteria | Kitasatospora sp. CB02891 | C20 | BG | 517 | Tpn2 | A0A2M9LDX2 |
| Bacteria | Amycolatopsis tolypomycina | C25 | BG | 534 | AtoE | WP_091316343.1 |
| Bacteria | Streptomyces showdoensis | C15 | BG | 533 | DMS1 | A0A2P2GK84 |
| Bacteria | Aquimarina spongiae | C15 | XB | 536 | DMS2 | WP_084549426.1 |
| Bacteria | Alicyclobacillus acidocaldarius | C30 | BG | 631 | SHC | WP_012811690.1 |
| Bacteria | Bacillus subtilis 168 | C30 | BG | 632 | SqhC | AFQ57872.1 |
| Bacteria | Streptomyces argenteolus | C40 | BG | 545 | Lon15 | BAF98632.1 |
| Bacteria | Methylococcus capsulatus Texas | C30 | BG | 670 | OSC | WP_154656699.1 |
| Bacteria | Rhodopirellula lusitana DSM 25457 | Unknown | BG | 664 | Unknown | WP_283431395.1 |
| Bacteria | Scytonema sp. PCC 10023 1680591 | C20 | B | 366 | MstE | ATN39899.1 |
| Bacteria | Enhygromyxa salina DSM 15201 | C30 | BG | 695 | OSC | WP_052558171.1 |
| Bacteria | Eudoraea adriatica | C30 | BG | 653 | EUS | WP_169336908.1 |
| Bacteria | Methyloceanibacter caenitepidi | C30 | BG | 662 | OSC | WP_045368728.1 |
| Bacteria | Methylococcus capsulatus Bath | C30 | BG | 670 | OSC | MCA2873 |
| Bacteria | Methylococcus capsulatus Bath | C30 | BG | 654 | SHC | MCA0812 |
| Bacteria | Methylomicrobium alcaliphilum 20Z | C30 | BG | 750 | OSC | MEALZ_0768 |
| Bacteria | Methylomicrobium alcaliphilum 20Z | C30 | BG | 655 | SHC-I | MEALZ_2523 |
| Bacteria | Methylomicrobium alcaliphilum 20Z | C30 | BG | 652 | SHC-II | MEALZ_3097 |
| Eukarya | Tetrahymena thermophila SB210 | C30 | BG | 655 | STC | XP_001026696.2 |
| Fungi | Phaeosphaeria sp. L487 | C20 | ABG | 946 | FCPS-KS | BAA22426.1 |
| Fungi | Gibberella fujikuroi | C20 | ABG | 952 | GfCPS-KS | Q9UVY5.1 |
| Fungi | Sphaceloma manihoticola | C20 | ABG | 962 | SmCPS-KS | CAP07655.1 |
| Plant | Abies grandis | C20 | ABG | 868 | AgAS | Q38710.1 |
| Plant | Arabidopsis thaliana | C20 | ABG | 802 | GA1 | AAA53632.1 |
| Plant | Picea glauca | C20 | ABG | 761 | PgCPS | ADB55707.1 |
| Plant | Physcomitrium patens | C20 | ABG | 881 | PpCPS-KS | BAF61135.1 |

| | | | | | | |
|---|---|---|---|---|---|---|
| Plant | Selaginella moellendorffii | C20 | ABG | 750 | SmCPS-KSL1 | AEK75338.1 |
| Plant | Arabidopsis thaliana | C20 | ABG | 877 | TPS04-GES | NP_564772.1 |
| Plant | Pisum sativum | C20 | ABG | 801 | CPS | AAB58822.1 |
| Plant | Stevia rebaudiana | C20 | ABG | 787 | Cpps1 | AAB87091.1 |
| Plant | Ginkgo biloba | C20 | ABG | 873 | GbTPS-Lev | AAL09965.1 |
| Plant | Picea abies | C20 | ABG | 867 | PaTPS-Iso | AAS47690.2 |
| Plant | Picea abies | C20 | ABG | 859 | PaTPS-LAS | AAS47691.1 |
| Plant | Hordeum vulgare subsp. vulgare | C20 | ABG | 826 | CPS | AAT49065.1 |
| Plant | Zea mays | C20 | ABG | 827 | CPS | AAT70083.1 |
| Plant | Pinus taeda | C20 | ABG | 850 | CPS | AAX07435.1 |
| Plant | Picea sitchensis | C20 | ABG | 761 | CPS | ADB55709.1 |
| Plant | Cistus creticus subsp. creticus | C20 | ABG | 808 | CPS | ADJ93862.1 |
| Plant | Lactuca sativa | C20 | ABG | 799 | CPS | BAB12440.1 |
| Plant | Oryza sativa Japonica Group | C20 | ABG | 773 | CPS | BAD42449.2 |
| Plant | Triticum aestivum | C20 | ABG | 797 | CPS | BAH56558.1 |
| Plant | Triticum aestivum | C20 | ABG | 831 | CPS | BAH56560.1 |
| Plant | Oryza sativa Japonica | C20 | ABG | 867 | CPS | NP_001403441.1 |
| Plant | Gossypium arboreum | C30 | BG | 742 | CAS | KHG12373.1 |
| Animal | Homo sapiens | C30 | BG | 732 | OSC | NP_001001438.1 |

**Supplementary Table 6:** Proteins that have been crystallized in the terpenoid cyclase family and the greater alpha-alpha toroid fold. PDB codes indicate crystal structures used for phylogenetic analysis in this study.

| Enzyme | Reaction type | Rxn class | Active site motif | Protein domains | Crystal organism | Domain of life | Crystal |
|---|---|---|---|---|---|---|---|
| isoprene synthase | isoprenoid diphosphate lyase | I | DDXXD, NSE | AB | *Populus canesscens* | euk | 3N0F |
| (+)-bornyl diphosphate synthase | monoterpene (C10) cyclase | I | DDXXD, NSE | AB | *Salvia officinalis* | euk | 1N1B |
| (-)-limonene synthase | monoterpene (C10) cyclase | I | DDXXD, NSE | AB | *Mentha spicata* | euk | 2ONG |
| (+)-limonene synthase | monoterpene (C10) cyclase | I | DDXXD, NSE | AB | *Citrus sinensis* | euk | 5UV0 |
| gamma-terpinene synthase | monoterpene (C10) cyclase | I | DDXXD, NSE | AB | *Thymus vulgaris* | euk | 5C05 |
| cineole synthase | monoterpene (C10) cyclase | I | DDXXD, NSE | AB | *Streptomyces clavuligerus* | bact | 2J5C |
| alpha-bisabolol synthase | sesquiterpene (C15) cyclase | I | DDXXD, NSE | AB | *Artemisia annua* | euk | 4FJQ |
| alpha-bisabolene synthase | sesquiterpene (C15) cyclase | I | DDXXD, NSE | ABG | *Abies grandis* | euk | 3SAE |
| epi-aristolochene synthase | sesquiterpene (C15) cyclase | I | DDXXD, NSE | AB | *Nicotiana tabacum* | euk | 5EAS |
| (+)-delta-cadinene synthase | sesquiterpene (C15) cyclase | I | DDXXD, NSE | AB | *Gossypium arboreum* | euk | 3G4D |
| drimenyl diphosphate synthsae | sesquiterpene (C15) cyclase | II | DXDD | BG | *Streptomyces showdoensis* | bact | 7XQ4 |
| taxadiene synthase | diterpene (C20) cyclase | I | DDXXD, NSE | ABG | *Taxus brevifolia* | euk | 3P5P |
| ent-copalyl synthase (plant) | diterpene (C20) cyclase | II | DXDD | ABG | *Abrabidopsis thaliana* | euk | 3PYA |
| ent-copalyl synthase (bact) | diterpene (C20) cyclase | II | DXDD | BG | *Streptomyces platensis* | bact | 5BP8 |
| tuberculosinyl diphosphate synthase | diterpene (C20) cyclase | II | DXD | BG | *Mycobacterium tuberculosis* | bact | 6VPT |
| terpentedienyl diphosphate synthase | diterpene (C20) cyclase | II | DXDD | BG | *Kitasatospora sp.* CB02891 | bact | 7XKX |
| squalene-hopene cyclase | triterpene (C30) cyclase | II | DXDD | BG | *Alicyclobacillus acidocaldarius* | bact | 2SQC |
| oxidosqualene cyclase | triterpene (C30) cyclase | II | DC | BG | *Homo sapiens* | BOTH | 1W6K |
| abietadiene synthase | bifunctional terpenoid cyclase | I/II | DDXXD, NTE, DXDD | ABG | *Abies grandis* | euk | 3S9V |
| linalool dehydratase isomerase | terpene isomerase | n/a | n/a | B | *Castellaniella defragrans* | bact | 5HLR |

| | | | | | | | |
|---|---|---|---|---|---|---|---|
| meroterpene cyclase | meroterpene cyclase | II | DxD | B | *Scytonema* | bact | 6SBB |
| protein farnesyltransferase | protein prenyltransferase | n/a | n/a | BX | *Cryptococcus neoformans (apo); R. norvegicus (complexed)* | euk | 1FPP |
| protein geranylgeranyl-transferase (type I) | protein prenyltransferase | n/a | n/a | BX | *Rattus norvegicus* | euk | 1N4P |
| Rab geranylgeranyl-transferase (type II) | protein prenyltransferase | n/a | n/a | BX | *Rattus norvegicus* | euk | 1DCE |
| Rab geranylgeranyl-transferase (type III) | Protein prenyltransferase | n/a | n/a | BXY | *Homo sapiens* | euk | 6J74 |

**Supplementary Table 7:** Functional regions for evolutionary rate shift analysis.

| Region | Definition (as residues of *Alicyclobacillus acidocaldarius* SHC; PDB ID 2SQC) | Graphical representation |
|---|---|---|
| N terminus | M1-D28 | 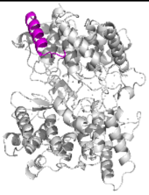 |
| C terminus | M611-R631 | 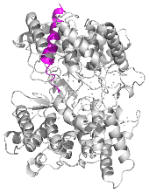 |
| Membrane anchor | W216-V233 | 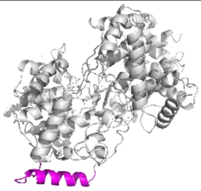 |
| Dimer contact surface | F236-D276 | 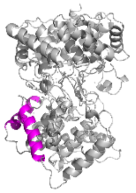 |
| Substrate tunnel | V128-F129, M147-P149, I152, L161, F166, A170, V174, L221, L225, I432, F434-F437, E438-V439 | 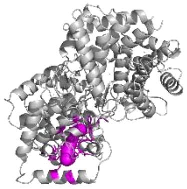 |
| Active site | L36, M42, I261-P263, A306-S307, W312, F365, D374, D376-D377, Y420, W489, G600-Y601, F605, L607, Y609, Y612 | 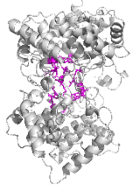 |
